# Recovery of genomic and transcriptomic profiles from decades-old FFPE brain tissues

**DOI:** 10.64898/2026.04.20.719637

**Authors:** Caroline Robinson Christiansen, Emily Hansen Firoozfard, Nikolay Oskolkov, M. Thomas P. Gilbert, Sarah S. T. Mak, Martin Wirenfeldt, Christina Kjær, Emilio Mármol-Sánchez

**Author notes:** Corresponding authors: Emilio Mármol-Sánchez, Christina Kjær. Shared first authorship (Caroline Robinson Christiansen, Emily Hansen Firoozfard). These authors contributed equally (Christina Kjær, Emilio Mármol-Sánchez).

## Abstract

Neurological, neurodegenerative, and psychiatric disorders impose substantial morbidity and disability worldwide, yet their molecular basis remains incompletely understood, in part due to limited access to human brain tissue. The Danish Brain Collection, comprising brains from individuals who lived in Danish psychiatric institutions from the 1940s to the 1980s, represents a unique but largely untapped resource for retrospective molecular investigation. Here, we assess the feasibility of extracting and sequencing DNA and RNA from decades-old FFPE brain tissue.

We systematically evaluate how extraction and library preparation strategies influence nucleic acid yield and quality, and show that RNA end-repair prior to library preparation substantially enhances transcript diversity, improving data quality from highly degraded samples. Despite extensive fragmentation, we recover biologically informative transcriptomic profiles, including protein-coding and microRNA expression profiles that retain tissue-specific signatures.

These results establish the Danish Brain Collection as a viable resource for genomic and transcriptomic analyses and demonstrate the broader potential of archival FFPE tissues for large-scale molecular studies.

## Introduction

Neurological, neurodegenerative, and psychiatric disorders affect over three billion individuals worldwide, representing one of the most significant public health challenges of our time [1]. These conditions diminish the quality of life of affected individuals and impose enormous economic burdens on families and healthcare systems [2]. Yet our ability to diagnose, predict disease progression, and effectively treat these disorders remains limited. Next-generation sequencing of brain-derived DNA and RNA has begun illuminating the molecular mechanisms underlying conditions like schizophrenia and major depressive disorder [3], Alzheimer’s disease and related dementias [4], or epilepsy [5], revealing disease-associated genetic variants, disrupted pathways, and potential therapeutic targets. Despite these advances, routine clinical protocols still predominantly rely on symptom-based assessment, which carries inherent limitations: symptoms overlap between disorders, early disease presents subtly, and subjective interpretation can lead to diagnostic uncertainty [6]. This introduces the risk of misdiagnosis leading to inappropriate treatment [7]. Moreover, most pathologies remain challenging to treat [8]. Therefore, the current gap between clinical intervention and patients’ needs stems from an incomplete molecular understanding of the pathological changes in the brain underlying these diseases.

A major issue faced by researchers is that large-scale molecular sequencing studies ideally require fresh tissue in which nucleic acids remain intact. Acquiring such samples is exceptionally difficult due to ethical considerations, logistical challenges, and donor scarcity [9], which hinders our ability to capture the full spectrum of disease heterogeneity. Formalin-fixed paraffin-embedded (FFPE) tissue archives represent a potential alternative. Over the decades, health institutions have accumulated vast FFPE repositories encompassing diverse patient populations, disease states, and historical periods.

One such resource is the Danish Brain Collection (DBC) (https://brain-collection.rsyd.dk/). Assembled between 1945 and 1982, this repository comprises a registered 9,479 brains from Danish psychiatric patients, including an estimated 50,000 FFPE tissue blocks representing multiple brain regions, and spanning widely different treatment eras dominated by divergent infectious disease landscapes and environmental exposures. It ranks among the world’s largest brain collections and encompasses an exceptionally broad diagnostic spectrum: dementia, major depression, schizophrenia, epilepsy, Huntington’s and Parkinson’s disease, among others. Examining molecular signatures across this nearly four-decade window offers unique opportunities to investigate whether brain disorder biology has remained constant or evolved through time, something that contemporary biobanked archives alone cannot easily reproduce. Despite its potential, the Danish Brain Collection remains largely unexplored using modern molecular techniques. No systematic efforts have applied high-throughput sequencing to these samples, likely due to ethics restrictions and concerns about their age and degradation status. Indeed, formaldehyde fixation is known to create extensive cross-links and fragmentation in biomolecules present in the tissues [10], severely compromising genetic material integrity. For sequencing, these issues translate into low yield, poor library complexity, and high error rates [11]. Even after optimization efforts [12–15], the molecular viability of decades-old FFPE brain tissues archived in pathology collections has remained largely uncharacterized, particularly where fixation protocols were not standardized.

Nevertheless, despite the extensive induced cross-linking and chemical damage created in tissues upon FFPE treatment [16], valuable molecular information can still be recovered [17–20], even from spatially localized areas of a given FFPE block [21–23]. Indeed, compelling evidence suggests that the main limiting factor affecting nucleic acid integrity is not the age of the FFPE sample per se, but rather the chemical conditions with which the original tissues were treated [16,24–28]. Moreover, recent methodological developments have enabled the recovery of genomic and transcriptomic data from specimens up to thousands of years old using specialized protocols handling extreme fragmentation and chemical modifications [29–34]. Therefore, the main question is not whether genetic material may survive over time, but whether molecular information remains sufficiently intact to yield robust biological insights.

In this pilot study, we aimed to determine whether informative high-throughput DNA and RNA sequencing data from a subset of decades-old FFPE brain samples archived in the DBC can be isolated, sequenced and analyzed, so as to reconstruct tissue-specific gene expression patterns from the original human brain tissues. To do so, we first worked on optimizing laboratory methods for DNA and RNA extraction and library preparation and then applied custom bioinformatic pipelines inspired by previous reports on historical archived tissues from museum collections [32].

## Methods

### Study design

We used human post-mortem brain tissue in the form of formalin-fixed paraffin-embedded (FFPE) tissue archived at The Danish Brain Collection located at University of Southern Denmark, Odense, Denmark, and Open Patient data Explorative Network (OPEN). The Danish Brain Collection is a biobank comprising a registered 9,479 brains, which were obtained during autopsies conducted at Danish state mental hospitals from 1945 to 1982 (https://brain-collection.rsyd.dk/). A total of 10 samples from 4 patients were included, representing the following anatomical brain regions: pons, thalamus, cerebellar vermis, and corpus striatum. An overview of the 10 FFPE blocks used and their tissue age and identity is shown in **Table 1**. The included patients were identified by searching for “epilepsy” in a database of the original patient diagnoses, and then choosing patients based on information from discharge letters stored on a GDPR-secured server, both provided by the Open Patient data Explorative Network (OPEN), OUH and Department of Clinical Research, University of Southern Denmark, Odense, Denmark. The specific pathological cohort is not the focus of the current study, which is instead intended to develop a methodological framework for the molecular study of aged FFPE blocks stored at the Danish Brain Collection.

**Table 1:**
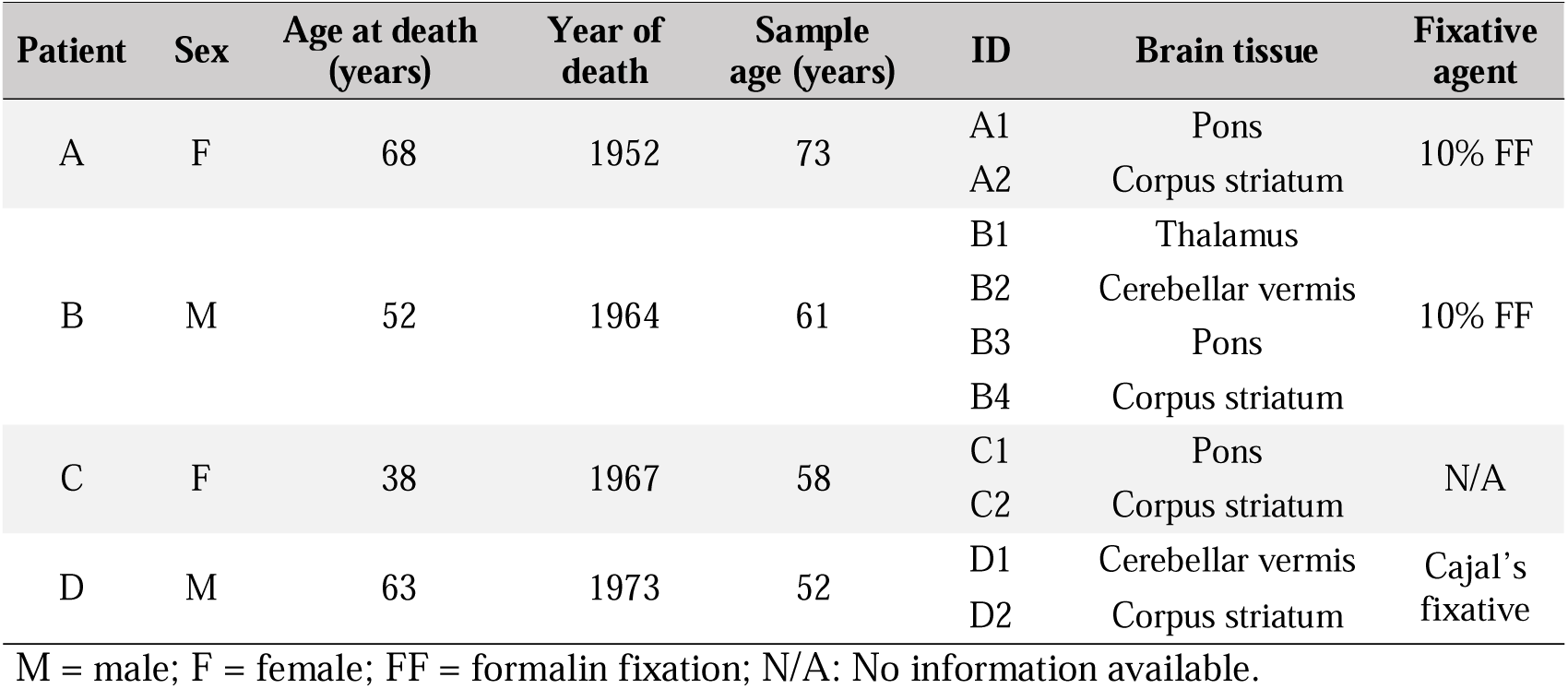
FFPE brain samples included in this study.

### Ethics declaration

This study was approved by The Danish National Committee on Health Research Ethics (ethical permission journal and supplement no. CVK-2308632 and CVK-112142) in accordance with the Danish Act on Research Ethics Review of Health Research Projects (Komitéloven) and all methods were performed in accordance with relevant guidelines and regulations. Archival human biological material used here was collected between 1945 and 1982, prior to the introduction of requirements for obtaining informed consent for the samples in question. Consequently, individual informed consent has not been obtained. This general issue concerning the DBC has been addressed previously in 2006 by The Danish National Committee on Biomedical Research Ethics journal number 2005-7049-6.

Under Danish law, research use of such archival material without consent may be permitted subject to approval by the competent research ethics committee, cf. Komitéloven. The present study has been reviewed and approved on this basis.

### Preprocessing

FFPE blocks from each patient were selected based on the registered time of death of the individual, to ensure that a range of different preservation ages were included in this pilot study to assess any potential effects of storage time on the quality of molecular data. Additionally, FFPE blocks containing tissue from four different neuron-rich brain areas were included, representing tissue-specific variation across different brain regions. Tissue identity of each block is provided in **Table 1**. First, each FFPE block was sectioned into three separate pieces to ensure comparable cell-rich grey matter content. To remove paraffin from the tissue, two pieces were incubated overnight at 50°C, then for 1 hour at 60°C. The first piece was left in its melted state (referred to as “Melted”) and stored at -20°C. The second piece was re-embedded in paraffin and not used any further in this study. The third piece remained unprocessed in its original state (referred to as “original FFPE”) and stored at room temperature in a dry, dark environment before being transferred to -20°C.

### Isolation of DNA and RNA

All laboratory procedures were carried out in clean laboratory facilities specifically dedicated to the processing of historical and/or ancient samples within the GLOBE Institute (University of Copenhagen, Denmark). DNA was extracted from FFPE blocks using three approaches: (i) the DNeasy Blood & Tissue Kit (Qiagen) with heat-mediated decrosslinking at 90°C [16], (ii) the same kit with an additional deparaffinization step using Deparaffinization Solution (Qiagen), and (iii) the AllPrep^®^ DNA/RNA FFPE Kit (Qiagen) for simultaneous DNA and RNA recovery. RNA was also independently extracted using the mirVana^TM^ microRNA Isolation Kit (Thermo Fisher Scientific), selected based on its prior successful application for RNA extraction from ancient tissue samples [32,34], with additional pretreatment steps including heat-mediated deparaffinization and decrosslinking. Detailed protocols for each extraction method, including reagent volumes, incubation conditions, and all modifications to the manufacturers’ recommendations, are provided in **Supplementary Methods**. Additional metadata for DNA and RNA extracts is also available in **Supplementary Tables 1** and **2**. Nucleic acid concentrations were measured using Qubit™ High Sensitivity dsDNA and RNA kits (Thermo Fisher Scientific) with a Qubit v3.0 fluorometer (Thermo Fisher Scientific). Size distribution and integrity of the DNA and RNA extracts for each sample were assessed using the High Sensitivity RNA ScreenTape, and High Sensitivity D1000 DNA ScreenTape in an Agilent 2200 TapeStation System (Agilent Technologies).

### Library preparation

DNA libraries were prepared following the Blunt-End-Single-Tube (BEST) protocol [35,36], comprising end-repair, adapter ligation, and fill-in reaction steps. Libraries were quantified by qPCR (Mx3005P, Agilent Technologies), indexed with unique dual 8-bp indices, and purified using HighPrep PCR beads (MagBio Genomics). cDNA libraries from RNA extracts (n = 20, from mirVana and AllPrep extractions) were prepared using the NEXTFLEX Small RNA-Seq Kit v4 with Unique Dual Indexes (Revvity), without size selection. For a subset of 10 RNA samples (from A1, A2, B2, C2, D1 pairs), end-repair was performed prior to additional library preparation using T4 Polynucleotide Kinase (T4 PNK, NEB) to repair damaged 5’ and 3’ termini on fragmented RNA molecules. A detailed protocol for all library preparation steps, including reagent compositions, thermocycler programs, and clean-up procedures, is provided in **Supplementary Methods**. Library concentrations were quantified using the Qubit™ dsDNA Quantification Assay Kit (Thermo Fisher Scientific) with a Qubit 2.0 Fluorometer (Thermo Fisher Scientific). The size distribution of the DNA and cDNA libraries was measured using the High Sensitivity DNA assay (Agilent Technologies) with a Bioanalyzer 2100 system (Agilent Technologies).

### Sequencing

The DNA (n = 33) and cDNA (n = 30) libraries were separately submitted to Novogene (Planegg, Germany), and independently sequenced on an Illumina NovaSeq X Plus system, with a full 10B lane in 150 bp paired-end configuration. For cDNA libraries, a homogeneous distribution of sequencing effort across all samples was applied. For DNA libraries, we balanced the sequencing effort so that each of the four original individuals obtained equivalent amounts of sequences at a target genome-wide coverage of at least 10×.

### Statistical analysis

Statistical comparison of DNA and RNA yield between extraction methods and pretreatment conditions was performed using two-sided paired t-tests in R v4.4.2.

### Sequence analysis

Raw FASTQ files were quality-filtered using fastp v0.24.1 [37] with the following parameters: automatic adapter detection and trimming, polyG tail removal, paired-end read merging, a minimum sequence length of 30 nt for DNA and 18 nt for RNA data, and a minimum overlap of 30 and 15 matching nucleotides for paired-end merging of DNA and RNA sequences, respectively. Only successfully merged paired sequences were retained for subsequent analyses. The human RefSeq assembly (GRCh38.p14, GCF_000001405.40) and associated gene annotations were used as reference. A reference transcriptome was generated using gffread v0.12.6 [38], retaining only the longest transcript isoform per gene across protein-coding messenger RNAs (mRNAs), ribosomal RNAs (rRNAs), transfer RNAs (tRNAs), long noncoding RNAs (lncRNAs), small nuclear RNAs (snRNAs), and small nucleolar RNAs (snoRNAs). The microRNA (miRNA) annotation was retrieved from miRGeneDB v3.0 [39]. Alignment was performed using Bowtie2 v2.4.2 [40] with end-to-end sensitive mode. DNA sequences were first sorted with the sambamba v0.8.0 tool [41], then deduplicated using the markdup function from SAMtools v.1.10 [42]. RNA sequences were not deduplicated, as the absence of unique molecular identifiers (UMIs) in the library preparation protocol prevents reliable distinction between PCR duplicates and genuinely identical RNA fragments derived from highly expressed loci. RNA library complexity was instead estimated as 1 minus the duplication rate reported by fastp. Genic and intergenic mapping was resolved using BEDtools v2.31.0 [43] and human gene coordinates. RNA sequences mapped to the reference transcriptome that were also identified as either overlapping intergenic or intronic regions genome-wide were flagged and removed. Taxonomic classification of DNA and RNA sequences was performed using KrakenUniq v0.7.3 [44] with a custom database [45] to characterize endogenous human content and exogenous contamination. Additional details are described in **Supplementary Methods**.

### Gene expression quantification

RNA sequences mapped to the human reference transcriptome were assigned to each corresponding gene with the intersect function from BEDtools v2.31.0 [43], and the percentage of the transcript body supported by any mapped read (breadth of coverage) was determined with the BEDtools coverage function. For miRNA genes, RNA sequences mapped to mature miRNA loci were considered as successful alignments if they overlapped the corresponding region by at least 75% of their length (bedtools intersect -f 0.75). Genes were considered successfully captured if they had a transcript-wide coverage >10%.

### Damage analysis

The PMDtools v0.60 [46] software was used to identify C>U deamination events in both RNA and DNA sequencing data. For DNA data, PCR-deduplicated sequences mapped to the GRCh38 assembly were analyzed over the first 50 nucleotides of each mapped sequence. For RNA, sequences mapped to the custom transcriptome were assessed over 30 nucleotides with single-stranded mode enabled. Only C>U deaminated sites, read as C>T by the sequencer, were considered.

### Clustering and tissue-specific gene expression analysis

Protein-coding mRNA and miRNA quantification for each RNA sample were used as input for Principal Component Analysis (PCA) after variance-stabilizing transformation (vst) with DESeq2 v1.44.0 [47]. For tissue-specific clustering, UMAP dimensionality reduction was applied using the umap v0.2.10.0 R package (https://github.com/tkonopka/umap). As modern reference, we used a comprehensive human RNA-seq atlas retrieved with the getAtlasExperiment function (E-MTAB-2836) from ExpressionAtlas v1.32.0 R package [48] including 11 healthy tissues [49,50], supplemented with chondrocyte samples [51] and four brain regions [52]. For the microRNA fraction, small RNA-seq datasets from human healthy tissues were used [52–54]. A comprehensive list of all reference samples is provided in **Supplementary Table 3**. Quantification values were normalized with the trimmed mean of M-values (TMM) method [55] using edgeR v4.2.1 [56] and log transformed. FFPE RNA data were then projected onto the learned UMAP embedding. To assess regional identity within different brain structures, end-repaired FFPE protein-coding mRNA profiles were projected onto a supplementary reference built from GTEx database v11 bulk RNA-seq gene read counts (GENCODE 47 annotation) obtained from GTEx portal (https://www.gtexportal.org), comprising cerebellar vermis, corpus striatum, cortex, hippocampus, and hypothalamus samples (**Supplementary Table 4**). End-repaired pons samples (A1) were excluded from these analyses since no pons tissue was available within the said GTEx reference expression atlas. Further details on reference data processing and UMAP parameters are provided in **Supplementary Methods**.

Hierarchical clustering was performed separately on protein-coding mRNA and miRNA profiles from end-repaired libraries (n = 10) using the top 100 most variable genes computed as those with gene-wise higher variance across log_2_-transformted CPM expression values from all end-repaired samples. TMM-normalized log -transformed CPM values were first converted to z-scores per gene and clustered with the Ward’s minimum variance method (ward.D2) and Euclidean distance, then visualized using ComplexHeatmap v2.20.0 [57]. Tissue specificity was quantified using the tau (τ) index [58], computed from the ExpressionAtlas reference multi-tissue atlas. The tau index was defined as:

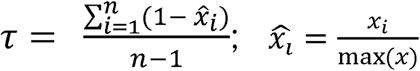

where n is the number of tissues, x□□ is the expression normalized to the tissue with maximum expression, and x□ is the average expression in tissue i. Tau values range from 0 (uniform expression) to 1 (tissue-restricted). Validated tissue markers were defined as genes with: (i) τ ≥ 0.5; (ii) peak expression in cerebellar vermis or corpus striatum in the reference atlas; (iii) detectable FFPE abundance (average log CPM ≥ 3); (iv) transcript coverage > 10% in the corresponding FFPE tissue (protein-coding genes only); and (v) concordant directionality between the reference atlas and FFPE data. Tissue identity was assessed through this marker-level framework rather than through differential expression against external brain datasets, since a formal cross-dataset differential-expression comparison is confounded in this setting by the absence of matched fresh-frozen controls and by platform, normalization, and region-definition differences between our FFPE bulk RNA-seq data and external references. In this way, the τ index is more robust to these differences, as it evaluates whether region-restricted expression is retained rather than comparing absolute expression levels across datasets. Full details on clustering and filtering criteria, as well as on gene selection are provided in **Supplementary Methods.**

### Gene set enrichment analysis

Gene ontology (Biological Process) over-representation analysis was performed separately on the validated sets of cerebellar vermis and corpus striatum marker genes using ShinyGO v0.85 [59], with all protein-coding genes evaluated for tissue specificity (n = 19,433) supplied as the custom background and an enrichment false discovery rate (FDR) threshold of 0.05. All remaining parameters were used as set by default except for the minimum pathway size, which was defined as a minimum of 5 genes.

## Results

We have analyzed post-mortem formalin-fixed paraffin-embedded (FFPE) brain tissue from the Danish Brain Collection, obtained from autopsies carried out at Danish state hospitals between 1945 and 1982 (**Table 1**). Both DNA and RNA were extracted in bulk, sequenced and analyzed comparing different laboratory approaches for pretreatment, nucleic acid isolation and transcriptional damage repair. Complete metadata information including pretreatment, extraction batch, sample identity, extraction method and nucleic acid yield is available in **Supplementary Table 1** for DNA and **Supplementary Table 2** for RNA extracts.

### Pretreatment of FFPE blocks and nucleic acid extraction method affect DNA yield

We first sought to compare the effectiveness of different methods for removing paraffin from FFPE brain tissues, and how this impacted the DNA concentration obtained after nucleic acid extraction. Overnight paraffin melting significantly increased DNA yield compared to original untreated FFPE samples without paraffin removal (*P*-value = 1.8 × 10^-4^), and this was also observed when applying a deparaffinization solution (**Fig. 1A**), although it did not reach statistical significance (*P*-value = 0.1752). All samples tested for pretreatment conditions and their DNA yield are in **Supplementary Table 5**.

**Figure 1:**
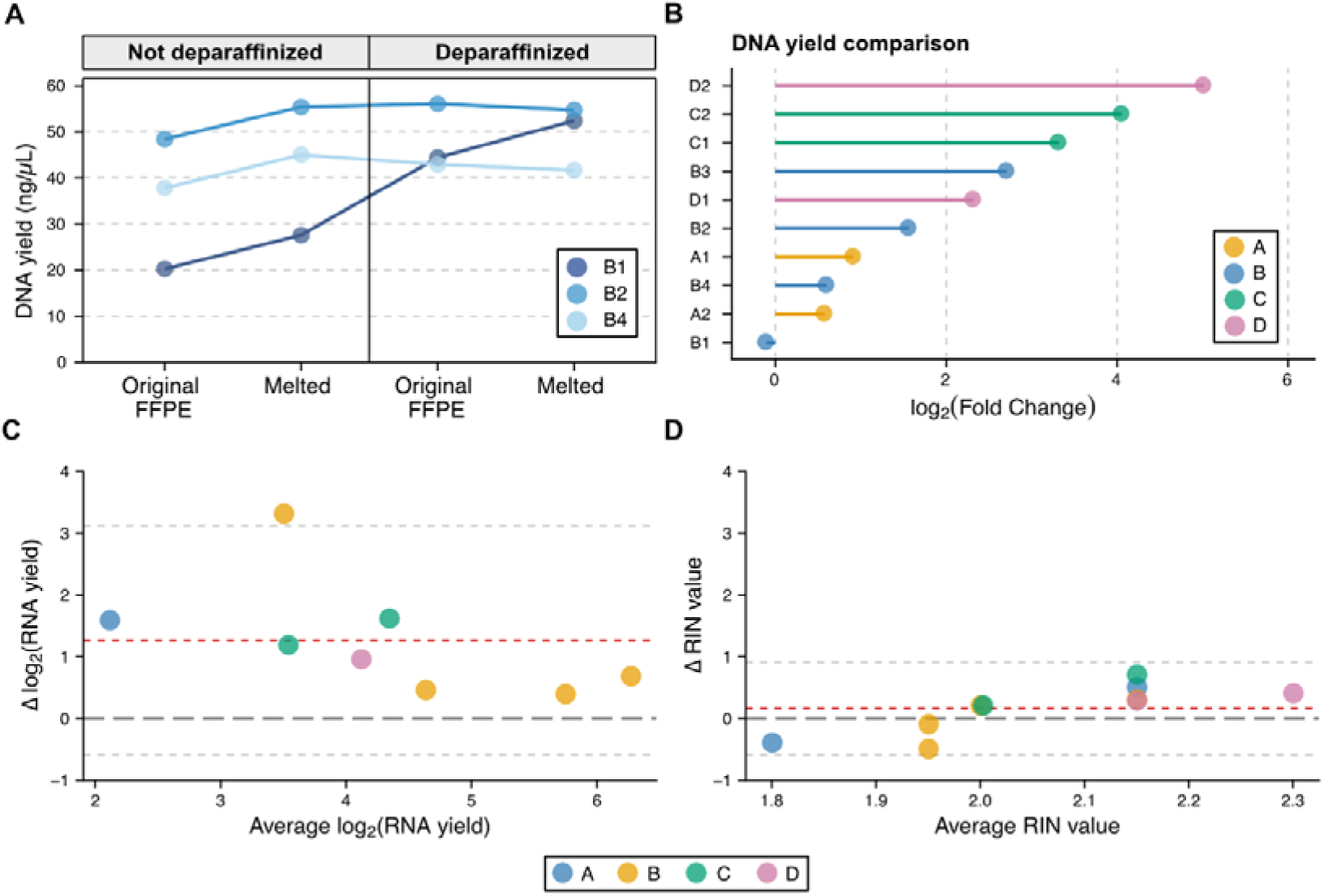
Pretreatment and extraction protocol impact on DNA and RNA yield. (**A**) DNA yield obtained from FFPE samples B1, B2, and B4 using two pretreatment approaches: physical removal of excess paraffin (original vs. melted samples) and enzymatic deparaffinization (see Methods). (**B**) Fold change of the difference in DNA yield between the AllPrep and Blood & Tissue methods for DNA extraction. A positive fold change value means a higher DNA yield was obtained with the AllPrep protocol, and vice versa. (**C**) Bland-Altman plot on the agreement between AllPrep and mirVana methods for RNA extraction according to the yield obtained with both protocols. The dark red dotted line represents the average difference in RNA yield in log_2_ scale. Light grey dotted lines represent the upper and lower confidence intervals at ±1.96 standard deviations from the average. (**D**) Bland-Altman plot on the agreement between AllPrep and mirVana methods for RNA extraction according to the RNA integrity (RIN) values obtained with both protocols. The dark red dotted line represents the average difference in RIN values. Light grey dotted lines represent the upper and lower confidence intervals at ±1.96 standard deviations from the average. Additional data from patients A, B, C, and D can be found in **Table 1**.

Next, we compared two different DNA extraction approaches on FFPE blocks. The DNeasy Blood & Tissue method from Qiagen is designed for simple and efficient recovery of DNA from different types of tissues. Incidentally, it can also recover RNA molecules during the extraction protocol [16], but we did not use this fraction from this particular kit. Moreover, it is not specifically tailored for FFPE samples, so we adapted it to include an initial decrosslinking step following prior reports demonstrating the efficacy of heat-mediated decrosslinking for FFPE DNA recovery [16]. In contrast, the other method we tested (AllPrep^®^ DNA/RNA FFPE, also from Qiagen) is specifically designed to work with FFPE tissues and allows simultaneous isolation of DNA and RNA molecules from the same sample. Despite this, it is a more time-consuming and involved protocol requiring more than double the time for its manual implementation. We again extracted DNA in an independent batch from all ten FFPE brain samples using both approaches. Upon comparison of both extraction kits, we found differences in the DNA yield distribution, with an overall higher but variable DNA concentration obtained with the AllPrep protocol compared to the Blood & Tissue method (**Fig. 1B**). Numeric values for total DNA concentration in all analyzed samples of this batch are available in **Supplementary Table 6**.

### Extraction protocol affects RNA yield but not RNA integrity

Next, to determine the efficacy of RNA isolation from decades-old FFPE tissues, we tested two different kits and evaluated both RNA yield and transcript integrity (as expressed by RIN values). We again compared the AllPrep kit, which allows RNA extraction jointly with DNA, and the mirVana^TM^ miRNA Isolation Kit (Thermo Fisher Scientific), which is optimized for short-length RNA recovery. The extraction method significantly influenced RNA yield, but not the RNA integrity. The use of the AllPrep kit resulted in extracts with significantly higher RNA concentration than the mirVana protocol (*P*-value = 3.79 × 10^-3^). Interestingly, the AllPrep method showed a higher tendency to improve the RNA yield over the mirVana kit at lower concentrations (**Fig. 1C**). Conversely, there were no significant differences in RIN values between the mirVana and AllPrep techniques (**Fig. 1D**, *P*-value = 0.2197), and the AllPrep kit only marginally improved the RNA integrity obtained at RIN values >2. Numeric values for total RNA concentration and RIN values in all analyzed samples are available in **Supplementary Table 7**.

### Decades-old FFPE brain samples yield mappable nucleic acids

After sequencing and mapping to the human (GRCh38) genome assembly, both DNA and RNA from FFPE brain tissues showed a high alignment rate, with overall above 75% of sequences successfully mapped (**Supplementary Table 8**, **Figs. 2A** and **2B**). However, DNA sequences belonging to patient D had ultra-low successful mapping, with only about 1.5% of them mapped to the human assembly (**Supplementary Table 8**, **Fig. 2A**). The remaining unmapped sequences in patient D libraries were attributable to a combination of microbial DNA consistent with environmental contamination (**Supplementary Table 9**, see Methods), library artifacts, and unmappable ultrashort fragments resulting from severe DNA degradation. These observations, combined with high duplication rates (**Supplementary Table 8**) indicate severe DNA degradation in these FFPE blocks, likely reflecting variation in historical fixation conditions within the collection as described in **Table 1**.

**Figure 2:**
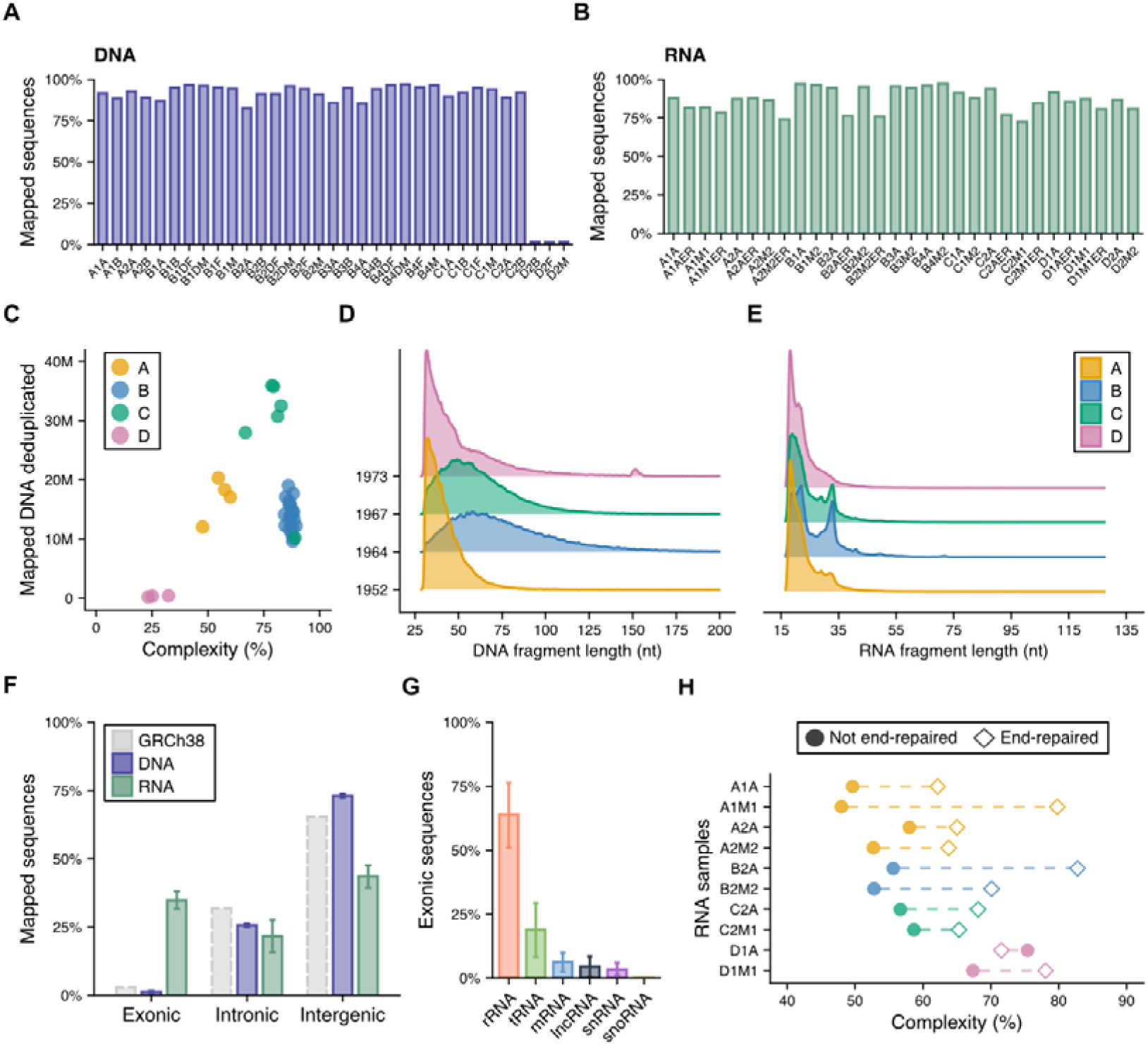
Sequencing and mapping statistics of DNA and RNA libraries from decades-old FFPE brain samples. (**A**) Percentage of DNA sequences successfully mapped to the human genome assembly (GRCh38) per library (see **Table 1** and **Supplementary Table 1** for additional data). (**B**) Percentage of RNA sequences successfully mapped to the human genome assembly (GRCh38) per library (see **Table 1** and **Supplementary Table 2** for additional data). (**C**) Library complexity of DNA samples, defined as the percentage of uniquely mapped sequences after deduplication. Fragment length density distribution of (**D**) DNA and (**E**) RNA sequences mapped to the human genome, grouped by patient and the recorded date of death highlighting the age of the FFPE blocks. (**F**) Proportion of mapped sequences overlapping exonic, intronic, and intergenic regions for DNA and RNA samples. Grey dashed values represent the proportion of exonic, intronic and intergenic regions annotated genome-wide in the human assembly (GRCh38). See Methods for more details on the loci considered. (**G**) RNA biotype composition of sequences mapped to exonic regions, showing the relative proportion of ribosomal RNA (rRNA), transfer RNA (tRNA), protein-coding messenger RNA (mRNA), long-noncoding RNA (lncRNA), small nuclear RNA (snRNA), and small nucleolar RNA (snoRNA) annotated loci. (**H**) Library complexity of RNA samples processed with and without T4 PNK end-repair prior to library preparation.

Consistent with this observation, samples from patients A, B, and C clearly outperformed those from patient D not only in the number of mapped DNA sequences, but also in library complexity (**Fig. 2C**), i.e. the number of uniquely aligned sequences after deduplication. Moreover, the fragment length distribution of successfully mapped DNA sequences was markedly shifted toward shorter fragments in samples from patients A and D compared to patients B and C, which showed longer and more dispersed fragment lengths with an average of ∼60 nucleotides (**Fig. 2D**). Interestingly, patient D, despite being the most recently deceased of the four individuals (date of death recorded in 1973, **Table 1**), showed shortened DNA fragments mapped to the human assembly, similar to the profiles observed for patient A, the oldest of the four (**Fig. 2D**). This, together with the very low number of mapped DNA sequences and extensive degradation of DNA in the FFPE blocks from this individual, suggests that chemical treatment during fixation, rather than age, is the main driver of DNA quality observed in our data.

These patterns contrast with those observed for the mapped RNA sequences, where most fragments were of ultrashort length, between 18 and 30 nucleotides (**Fig. 2E**). As with DNA samples, no clear age-related differences in RNA fragment length distribution were observed. Interestingly, contrary to what was observed in DNA, FFPE RNA from patient D samples showed profiles similar to the rest (**Supplementary Table 8**), including scarce microbial RNA contamination (**Supplementary Table 10**). Nevertheless, despite their short overall fragment length, RNA sequences showed clear signs of transcriptome-like mapping patterns, with nearly 35% of all mapped fragments genome-wide falling within annotated exonic regions of the human genome assembly (GRCh38, **Fig. 2F, Supplementary Table 8**). Intronic and intergenic mapping RNAs accounted for approximately 22% and 44% of the sequences, respectively. Conversely, DNA sequences showed markedly different proportions, more closely reflecting the overall genomic distribution of exonic, intronic, and intergenic regions, where DNA fragments mapped to exonic regions accounted for less than 2% of the total mapped sequences (**Fig. 2F**). Among exonic RNA sequences, the vast majority overlapped with rRNA loci, followed by tRNA, and protein-coding loci (**Fig. 2G**). Other less abundant noncoding loci such as lncRNAs, snRNAs and snoRNAs were also detected (**Fig. 2G**). Comprehensive abundance and transcript coverage profiles for all transcript types analyzed (see Methods) are available in **Supplementary Tables 11**-**16.**

### End-repair treatment increases library complexity but shortens RNA fragments

We next evaluated the effect of end-repairing damaged RNA fragment termini from decades-old FFPE samples on adapter ligation efficiency during library preparation, and its impact on the resulting transcriptomic sequencing data. To do so, we processed 10 samples (**Supplementary Table 2**) with and without applying T4 PNK end-repair prior to library preparation (see Methods). All end-repaired samples except for one (D1A) showed a relative increase in library complexity (**Fig. 2H**), defined as the fraction of non-redundant (i.e. non-PCR-duplicate) RNA fragments. Samples without end-repair treatment had an average 57% complexity, meaning that over 40% of sequencing effort likely consisted of redundant PCR duplicates, while end-repaired samples showed an average 71% complexity (**Fig. 2H**).

Consistent with the higher library complexity observed for end-repaired RNA samples, such treatment also significantly increased the number of protein-coding mRNA loci profiled at >10% transcript coverage (**Figs. 3A** and **3B**), a threshold we used as a heuristic for reliably detected genes [32]. This effect was observed across all 10 samples evaluated with and without end-repair treatment (**Supplementary Figs. 1**-**2**). Furthermore, for some of the more degraded and poorly profiled samples, particularly those from patients A and C, end-repair was essential to recover any protein-coding mRNA genes above this coverage threshold (**Supplementary Figs. 1**-**2**). Interestingly, several highly abundant mRNA loci detected in samples without end-repair (**Fig. 3A**), although at ultra-low coverage (<10%), showed reduced abundance but increased transcript coverage after end-repair (**Supplementary Table 11** and **Fig. 3B**).

**Figure 3:**
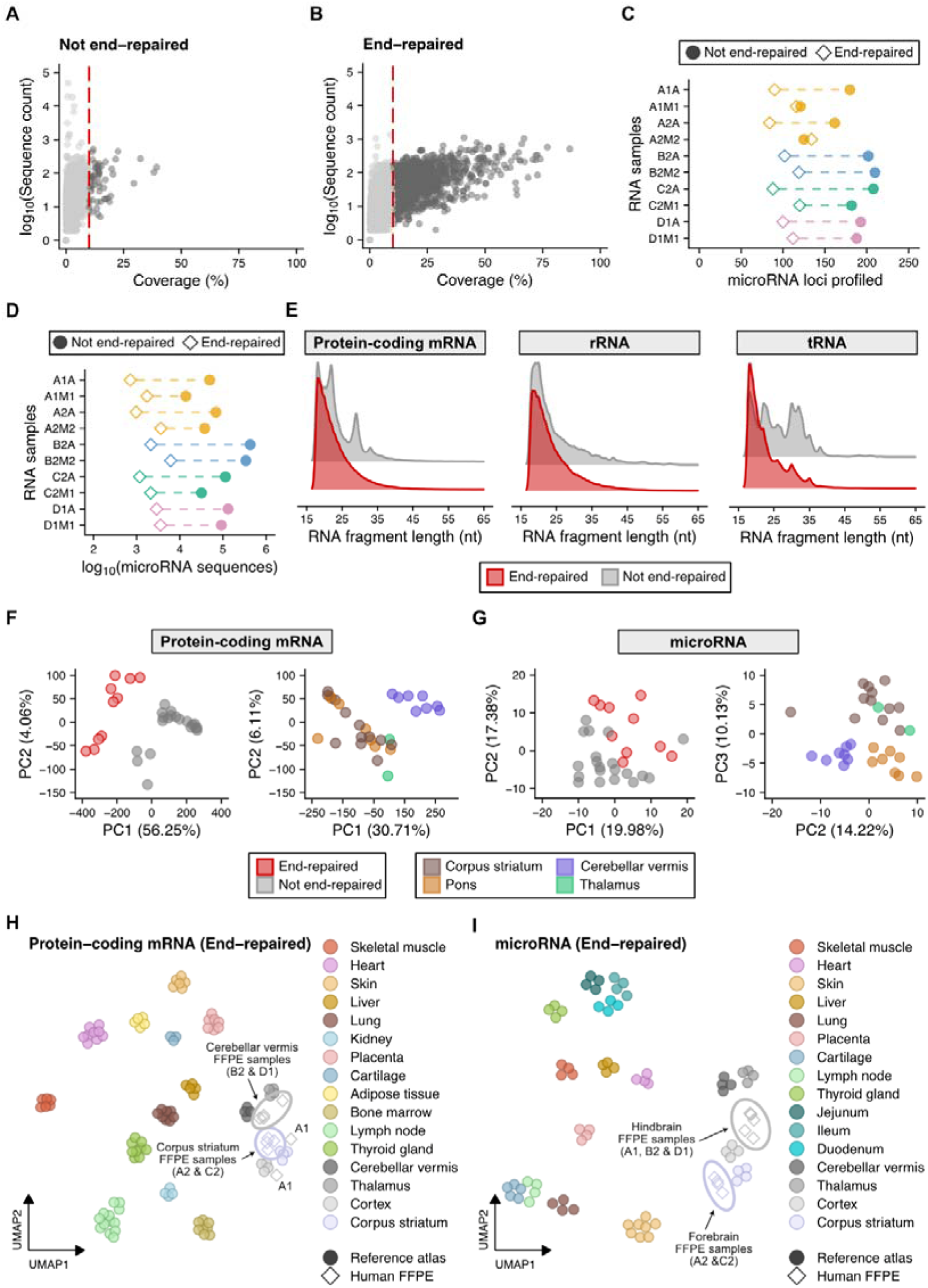
End-repair effects on RNA library composition and tissue-specific gene expression profiling from FFPE brain samples. (**A**) Sequence count vs. transcript coverage for protein-coding mRNAs in the 2A library from patient B without end-repair treatment (B2A sample). The red dashed line marks a 10% coverage threshold, defined as the lower limit to consider a given protein-coding mRNA gene as successfully profiled. (**B**) Sequence count vs. transcript coverage for protein-coding mRNAs in the 2A library from patient B with end-repair treatment (B2AER sample). (**C**) Number of microRNA loci profiled per sample, comparing libraries with and without end-repair treatment. (**D**) Number of mapped microRNA sequences per sample, comparing libraries with and without end-repair treatment. (**E**) RNA fragment length density distribution for protein-coding mRNA, rRNA, and tRNA genes, comparing end-repaired (red) and non-end-repaired (grey) libraries. (**F**) PCA of protein-coding mRNA expression profiles across all 30 FFPE brain samples. The left panel shows all samples colored by end-repair status. The right panel shows the same data after regressing out the end-repair effect, with samples colored by brain region. (**G**) PCA of microRNA expression profiles across all 30 FFPE brain samples. The left panel shows all samples colored by end-repair status. The right panel shows the same data with samples colored by brain region. (**H**) UMAP embedding of protein-coding mRNA expression from end-repaired FFPE brain samples (n = 10) projected onto a human multi-tissue RNAseq reference atlas. (**I**) UMAP embedding of microRNA expression from end-repaired FFPE brain samples projected onto a human multi-tissue small RNAseq reference atlas. FFPE sample pairs per tissue belong to patients A (n = 4, A1: Pons & A2: Corpus striatum), B (n = 2, B2: Cerebellar vermis), C (n = 2, C2: Corpus striatum), and D (n = 2, D1: Cerebellar vermis). The hindbrain region comprises pons and cerebellar vermis tissues, while the forebrain region comprises corpus striatum tissue.

This pattern suggests that end-repair enables a more even sampling of mRNA fragments across transcripts, rather than libraries being dominated by PCR duplicates of a small number of highly abundant loci. However, the increased resolution obtained for the protein-coding fraction in end-repaired RNA libraries did not extend to microRNA loci. Libraries without end-repair treatment yielded a higher number of profiled microRNA genes (**Fig. 3C**) and more mapped RNA fragments overlapping miRNA loci compared to end-repaired samples (**Fig. 3D**), suggesting that end-repair is detrimental for microRNA recovery. A comprehensive quantification of all microRNAs detected is available in **Supplementary Table 17**.

Besides, end-repaired samples also showed a reduced average RNA fragment length compared to samples without end-repair treatment, an effect that was consistent across all gene types analyzed but particularly pronounced for the most abundant transcript classes: rRNAs, tRNAs, and protein-coding mRNAs (**Fig. 3E**).

### Transcript length affects per-gene recovery in FFPE libraries

We explored whether intrinsic gene features are associated to transcript recovery. To do so, we examined per-gene transcript coverage and abundance as a function of transcript length and GC content across all detected protein-coding genes (coverage >10% and at least 1 count per million in log_10_ scale; **Supplementary Fig. 3**) in the ten end-repaired RNA libraries. Transcript length was the feature most associated with recovery, correlating positively but shallowly with per-gene abundance (Spearman *ρ* = 0.33). This agrees with longer transcripts accumulating more mapped sequences while being supported over a smaller proportion of their length. Conversely, GC content showed little to no association with either transcript coverage or abundance.

### Damage profiles indicate moderate cytosine deamination in FFPE-derived DNA and RNA

To assess the extent of chemical damage in the sequenced nucleic acids, we examined nucleotide misincorporation patterns along mapped DNA and RNA fragments. For DNA, all samples showed a characteristic cytosine-to-uracil (C>U) mismatch enrichment at the 5’ end of fragments and the complementary guanine-to-adenine (G>A) enrichment at the 3’ end (**Supplementary Fig. 4**), consistent with cytosine deamination induced by formalin fixation and post-mortem hydrolytic damage, as typically observed in degraded and archival DNA [10,60,61]. The magnitude of these terminal misincorporation frequencies was moderate, reaching approximately 2-4% at the first position across all samples, with a gradual decay toward background levels within the first 5-10 nucleotides. This pattern was consistent across all four patients regardless of sample age. For RNA samples, cytosine deamination frequencies were assessed along end-repaired fragments (**Supplementary Fig. 5**). Average deamination rates ranged from less than 1% to ∼3.5%, with a slight enrichment at the 5’ terminal positions relative to the fragment body. Compared to the DNA profiles, RNA deamination rates were of similar or slightly higher magnitude but showed a more uniform distribution along the whole fragment length rather than a steep terminal enrichment, which is consistent with the combined effects of formalin-induced and hydrolytic cytosine deamination acting across the full length of single-stranded RNA molecules. Samples from patients A and D showed higher deamination rates compared to those from patients B and C (**Supplementary Fig. 5**), broadly consistent with the increased DNA fragmentation observed for these individuals (**Fig. 2D**).

### Decades-old FFPE brain samples retain tissue-specific gene expression signatures

To determine whether the transcriptomic data recovered from FFPE brain samples captured biologically meaningful gene expression patterns, we performed dimensionality reduction analyses on both protein-coding mRNA and microRNA quantification profiles. PCA on protein-coding mRNA expression confirmed that end-repair status was the dominant source of variance, accounting for 56.25% of the total variation along PC1 (**Fig. 3F**, left panel). After regressing out the end-repair effect, the corrected PCA (**Fig. 3F**, right panel) revealed a clear separation by brain region along the first two components, with cerebellar vermis samples clustering distinctly from pons, corpus striatum, and thalamus samples. For microRNAs, PCA showed a less structured pattern (**Fig. 3G**, left panel) when examining the end-repair status, but separation by brain region became apparent when examining PC2 vs. PC3 (**Fig. 3G**, right panel), again with cerebellar vermis as the most distinct tissue. No clear influence of sex was observed in the PCA clustering for either protein-coding mRNAs or microRNAs when considered at transcriptome-wide level altogether (**Supplementary Figs. 6A** and **6B**). To dissect whether sex-specific transcriptional differences were still preserved irrespective of PCA results, we further examined the expression of canonical sex-linked marker genes across all FFPE RNA libraries. The X-inactivation lncRNA *XIST* was detected in 11 out of 14 libraries from the two female donors (patients A and C) but only 2 out of 16 libraries from the two male donors (patients B and D). Conversely, nine Y chromosome-linked genes selected with no close X-linked equivalent (*RPS4Y1*, *RPS4Y2*, *DDX3Y*, *EIF1AY*, *UTY*, *ZFY*, *KDM5D*, *USP9Y*, and *NLGN4Y*) were consistently detected in male-donor libraries, ranging from 8 to 12 out of the 16 male libraries depending on the gene, compared to 0 to 6 out of the 14 female libraries. The residual female-related detection might be attributable to cross-mapping with X-linked more distant paralogs (**Supplementary Fig. 6C, Supplementary Table 18**). This sex-specific expression pattern, recovered consistently across extraction kits and end-repair conditions, shows that decades-old FFPE-derived transcriptomic profiles retain genuine sex-linked biological signal beyond what transcriptome-wide clustering alone is able to deconvolute (**Supplementary Figs. 6A-B**).

To place these expression profiles in a broader biological context, we first projected the FFPE brain protein-coding mRNA profiles from end-repaired samples onto a human multi-tissue reference atlas (**Fig. 3H**). The FFPE brain samples consistently clustered adjacent to reference brain tissue samples, distinct from all other tissues, including skeletal muscle, heart, liver, lung, kidney, skin, and gut-derived tissues, among others. Moreover, we found a clear divergence between end-repaired FFPE samples obtained from cerebellar vermis tissue compared to those from the corpus striatum (**Fig. 3H**). Samples from patient A (Pons tissue) did not cluster with other hindbrain-derived tissues such as the cerebellar vermis, nor did they form a separate cluster. This result shows that despite decades of formalin fixation and paraffin embedding, these archival brain samples retain gene expression signatures that are recognizably brain-specific, reflecting differentiated protein-coding transcriptional programs between forebrain and hindbrain regions in the central nervous system. A similar pattern was found when projecting FFPE samples without end-repair treatment onto the same multi-tissue protein-coding human reference atlas, with all samples clustering nearby or adjacent to brain-derived tissues (**Supplementary Fig. 6D**). However, only those obtained from cerebellar vermis tissue showed a clear tissue-specific clustering pattern.

For microRNA profiles, projection onto a multi-tissue human microRNA expression atlas also showed a clearly defined clustering pattern, with end-repaired FFPE brain samples dividing into those derived from the hindbrain (pons and cerebellar vermis), and those derived from the forebrain (corpus striatum), again reflecting a shared post-transcriptional regulatory program among brain tissues of common developmental origin (**Fig. 3I**). When considering FFPE samples without end-repair treatment, however, such division was not apparent, although all of them clustered adjacent to brain-derived tissues included in the microRNA expression reference (**Supplementary Fig. 6E**).

We also evaluated tissue regional specificity against a brain-focused reference panel (**Supplementary Table 4**). After projecting the end-repaired FFPE mRNA libraries, excluding pons samples (A1), cerebellar vermis libraries from patients B and D (B2 and D1) co-localized with the reference cerebellar vermis cluster (**Fig. 4A**). For corpus striatum libraries, sample A2M2ER was projected within the reference striatal cluster, whereas A2AER localized with the hypothalamus cluster, and both C2 libraries were projected within the cortex cluster. The displacement of A2AER might be consistent with low library complexity (**Supplementary Fig. 1**), as this library recovered the lowest number of reliably detected protein-coding genes among the end-repaired set (249 nuclear protein-coding genes with >10% coverage, versus 637–4,615 for the remaining end-repaired brain libraries; **Supplementary Table 8**), leaving limited tissue-specific signal for a transcriptome-wide projection. We then restricted the embedding to tissue-specific genes (τ ≥ 0.5, including those identified in cortex, cerebellar vermis, or corpus striatum tissues; **Supplementary Table 19** and **Fig. 4B**). Under this marker-restricted projection, A2AER relocated from the hypothalamus cluster to the corpus striatum cluster. In contrast, both C2 libraries continued to localize within the cortex cluster, while cerebellar vermis libraries remained colocalizing with their corresponding tissue identity.

**Figure 4:**
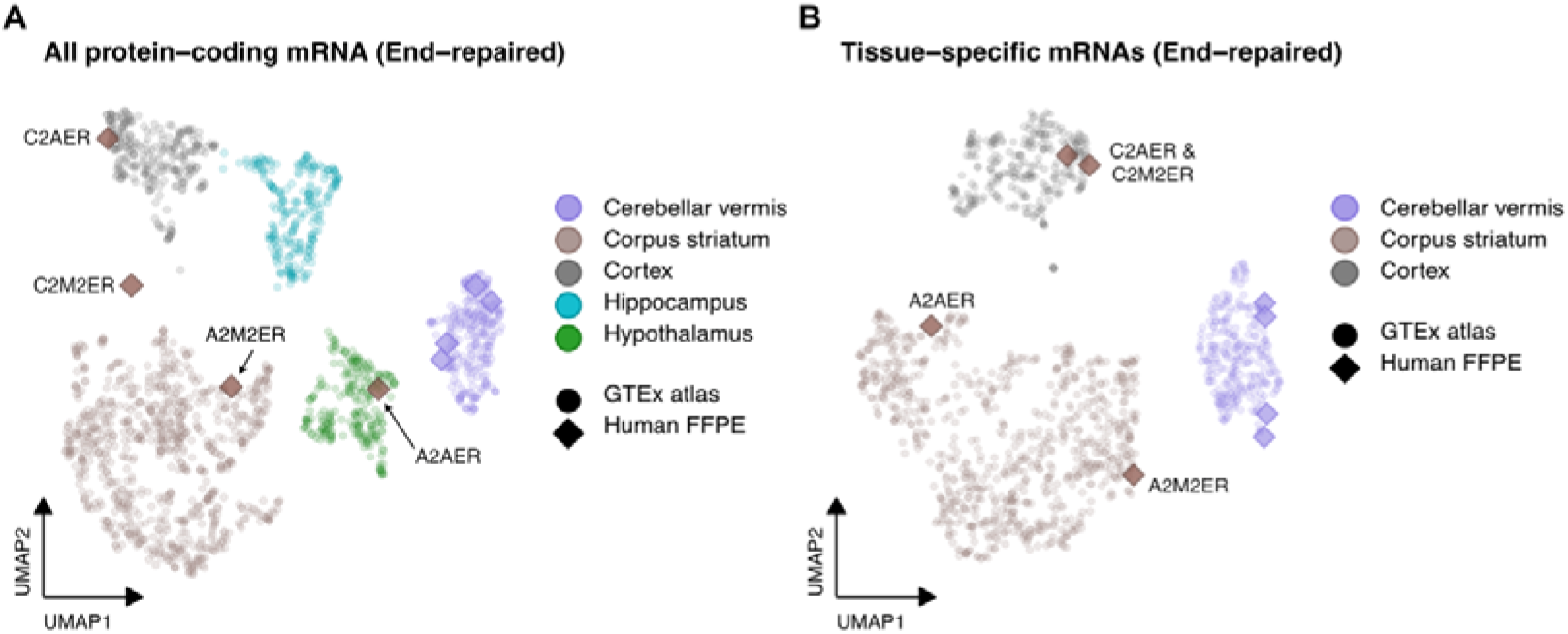
Regional brain specificity of end-repaired FFPE samples. (**A**) UMAP embedding of protein-coding mRNA expression from end-repaired FFPE brain samples projected onto a GTEx human brain reference atlas comprising cerebellar vermis, corpus striatum, cortex, hippocampus, and hypothalamus samples, using all annotated protein-coding genes. (B) UMAP embedding of the same libraries projected onto the brain reference atlas restricted to tissue-specific genes (τ ≥ 0.5 in cortex, cerebellar vermis, or corpus striatum; see Methods and **Supplementary Table 19**). FFPE libraries correspond to patients B (B2: cerebellar vermis) and D (D1: cerebellar vermis), and patients A (A2: corpus striatum) and C (C2: corpus striatum). Tissue identity of each block is provided in **Table 1**.

Furthermore, unsupervised hierarchical clustering of the top 100 most variable protein-coding mRNA genes separated the 10 FFPE samples primarily by patient and then by brain region, with cerebellar vermis and corpus striatum samples forming two distinct clusters (**Supplementary Fig. 7**). The two pons samples, both derived from patient A, clustered separately and were not further evaluated as region-specific. A similar tissue-driven separation was observed for the top 100 most variable miRNAs (**Supplementary Fig. 8**).

To determine whether this clustering reflects known regional gene expression patterns, we computed tissue specificity values (τ index) for all protein-coding mRNA and microRNA genes included in the multi-tissue gene expression collection used as reference (**Supplementary Tables 19**-**20**). Filtering criteria included tissue-restricted expression (τ ≥ 0.5), concordant directionality between the reference atlas and FFPE end-repaired samples, detectable abundance in FFPE samples, and adequate transcriptional coverage (see Methods). Across the full transcriptome, 321 protein-coding genes were validated as tissue markers: 177 in cerebellar vermis and 144 in corpus striatum. Of these, 28 fell within the top 100 most variable genes in the hierarchical clustering (18 cerebellar vermis, and 10 corpus striatum; **Supplementary Table 21**). Among the validated cerebellar vermis markers, several canonical cerebellar genes were identified, including the granule cell-associated zinc finger transcription factors *ZIC1* and *ZIC4*, the Purkinje cell synapse organizers *CBLN1* and *CBLN3*, the metabotropic glutamate receptor *GRM1,* the ETS transcription factor *ETV1*, and the voltage-gated potassium channel *KCNA1*, among others. Additional well-established cerebellar markers such as *GABRA6*, *PVALB* and *PAX6* were validated outside the top 100 variable gene set (**Supplementary Table 21**). Corpus striatum-enriched markers included the neuropeptide *CCK*, the postsynaptic calmodulin-binding protein *NRGN*, the calcium/calmodulin-dependent kinase *CAMK2A*, the synaptic protein *DDN*, and the cortexin *CTXN1*. The neuropeptides *SST* and *NPY* were among the strongest striatal markers validated outside the top 100 (**Supplementary Table 21**).

Gene ontology over-representation analysis of the validated protein-coding markers (**Supplementary Table 21**) confirmed their neural identity and showed a regional difference with functional emphasis: Cerebellar vermis gene markers (n = 177) were most significantly enriched for nervous system development (GO:0007399), synapse organization (GO:0050808), and neuron differentiation (**Supplementary Table 22**). In contrast, corpus striatum gene markers (n = 144) were most significantly enriched for synaptic signaling and neurotransmitter-release processes (**Supplementary Table 23**), including the synaptic vesicle cycle (GO:0099504), neurotransmitter secretion (GO:0007269), and synaptic vesicle exocytosis (GO:0016079). Both sets shared enrichment for core synaptic and neurodevelopmental processes, as expected for brain-derived tissue.

For miRNAs, 14 were validated as tissue-specific markers in end-repaired FFPE samples, of which 13 were among the 100 most variable according to hierarchical clustering (**Supplementary Table 24**). Four were detected as predominant in cerebellar vermis and ten in corpus striatum. The strongest cerebellar miRNA markers were miR-124 paralogues (miR-124-P1-v1 and miR-124-P2-v2), along with miR-376-P2 and miR-154-P14. Corpus striatum-related miRNAs included miR-137 (τ = 0.97), as well as miR-138-P2, miR-132, and miR-433 (**Supplementary Table 24**).

## Discussion

In this study, we demonstrate that decades-old FFPE brain tissue from the Danish Brain Collection retains sufficient molecular integrity for informative genomic and transcriptomic analyses. Systematic evaluation of extraction, library preparation, and bioinformatic approaches show that archival samples yield recoverable DNA and RNA with sequencing data capturing biologically meaningful, tissue-specific expression patterns. A summarized overview of our findings and recommendations is available at **Table 2**.

**Table 2:**
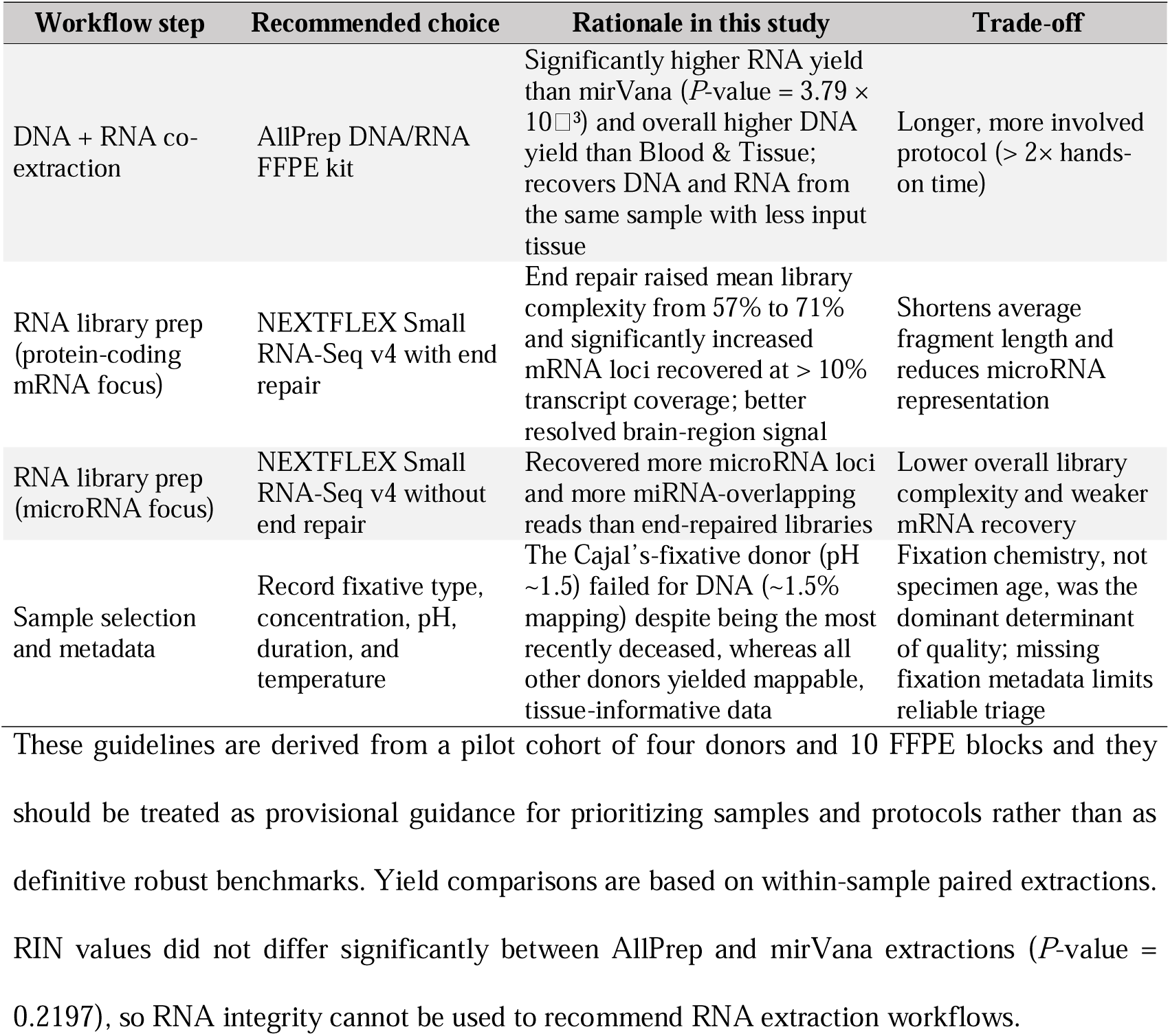
Recommended workflow for nucleic acid recovery from decades-old FFPE brain tissue, based on this pilot study.

### Practical considerations for nucleic acid recovery from aged FFPE tissues

Comparison of extraction methods identified the AllPrep protocol as the most suitable for DBC material, yielding higher DNA and RNA concentrations than Blood & Tissue and mirVana while requiring less input tissue (**Table 2**). Its ability to simultaneously recover DNA and RNA from the same specimen is particularly advantageous for limited archival material and enables parallel genomic and transcriptomic analyses.

We note, however, that the AllPrep protocol for DNA extraction is a significantly more involved workflow compared to the Blood & Tissue protocol, requiring about 2-fold more time to completion.

### Fixation chemistry drives variability in nucleic acid preservation

Not all samples performed comparably after nucleic acid extraction. Tissues from patient D exhibited markedly reduced DNA mapping rates and library complexity relative to the rest of the cohort. Notably, these samples were preserved using Cajal’s fixative rather than neutral buffered formalin. Cajal’s fixative is highly acidic (pH ∼1.5) [62], a condition known to accelerate depurination and phosphodiester bond cleavage in nucleic acids [25]. Consistent with prior reports, acidic or unbuffered fixation leads to increased fragmentation and reduced amplifiability compared to neutral conditions [16,28]. Therefore, the poor performance of patient D samples might be explained by the fixation chemistry rather than by tissue age, given that patient D is paradoxically the most recently deceased individual in the cohort (date of death 1973).

Patient A, the oldest individual in the cohort (1952), exhibited the shortest DNA fragment lengths and reduced RNA library complexity. The two available pons samples clustered with other brain regions. Nonetheless, they were probably insufficient (n = 2, single individual) for robust assessment of region-specific gene expression. Whether this reflects intrinsic properties of the pons, patient-specific degradation, or both cannot be resolved with the current data. Together with the findings from patient D, these results indicate that fixation chemistry, rather than specimen age, is the dominant determinant of nucleic acid preservation in archival material. Accordingly, detailed metadata on fixation conditions (i.e. fixative type, concentration, pH, duration, and temperature) should be systematically recorded and incorporated into study design, sample selection, and downstream analyses when working with FFPE collections (**Table 2**). For the Danish Brain Collection specifically, such information is sparse and only partially recoverable, which limits retrospective stratification of blocks by expected molecular quality and precludes a reliable collection-wide estimate of the fraction of samples suitable for sequencing. In the absence of complete fixation records, we suggest that suitability is better assessed empirically than predicted from archival metadata: a low-cost molecular pre-screen, such as shallow DNA sequencing to evaluate human mapping rate and fragment-length distribution, might reliably identify poorly preserved blocks (as exemplified by the acidic-fixative used in the sample from patient D, with ∼1.5% mapping) before committing full sequencing efforts. Where fixation records are accessible, neutral buffered formalin should be prioritized over acidic or unbuffered fixatives, which our data and prior reports associate with severe DNA degradation and reduced recoverability [16,25,28].

### End-repair improves FFPE transcriptomics

The application of T4 PNK end-repair prior to library preparation emerged as a key step for recovering informative transcriptomic data from highly degraded FFPE RNA. End-repaired libraries showed substantially higher complexity, with a greater proportion of sequences representing independent RNA molecules rather than PCR duplicates (**Table 2**). This improvement translated into enhanced gene detection sensitivity, enabling the tissue-specific gene expression profiling reported here.

The effect of T4 PNK end-repair likely reflects restoration of damaged 5’ and 3’ termini on fragmented RNA, which in their unrepaired state cannot efficiently ligate to sequencing adapters during library preparation. This is consistent with reports demonstrating improved mRNA capture from extracellular vesicles and plasma RNA [63,64], and extends these findings to the more severely degraded context of decades-old FFPE tissue. A trade-off is a relative reduction in microRNA representation, as the increased capture of longer repairable RNA fragments likely dilutes the small RNA fraction in the final library. This is an important consideration for study design, as omitting end-repair may be preferable when small RNAs with low relative abundance compared to other transcript types, such as microRNAs, are the primary target. Nevertheless, end-repaired libraries retained biologically informative miRNA profiles that recapitulated tissue-specific patterns observed in untreated samples, and more clearly resolved the hindbrain-forebrain distinction, consistent with differences in embryological origin and post-transcriptional regulation.

Furthermore, the shorter fragment length distribution observed in end-repaired libraries likely reflects either preferential restoration of termini on shorter damaged fragments, rendering them ligatable and thereby shifting the overall distribution, additional fragmentation introduced during enzymatic treatment and heat inactivation, or a combination of both (**Table 2**). Future optimization efforts could explore alternative inactivation strategies, such as bead-based cleanup without heat exposure, to preserve ultrashort fragments while retaining the gains in library complexity conferred by end-repair.

Transcript length was the main gene feature associated with per-gene recovery across the end-repaired libraries, i.e. longer transcripts accumulated more successfully mapped RNA sequences but were covered over a smaller proportion of their length, although such a correlation was not strong. We note, however, that this is a within-FFPE description of recovery among detected genes rather than a comparison against matched fresh-frozen material, which was not available. A predictive model of transcript preservation and recovery would require such matched controls and a larger cohort, and thus remains an avenue for future work.

### Biological validity of FFPE-derived transcript abundance profiles

A central finding of this study is that archival FFPE brain tissue retains recognizably tissue-specific transcriptomic signatures despite decades of formalin fixation and paraffin embedding. When projected onto independent modern multi-tissue reference atlases, FFPE brain samples clustered with brain-derived tissues and were clearly separated from non-neural tissues, across both protein-coding mRNA and microRNA profiles. Within the brain compartment, cerebellar vermis and corpus striatum were readily distinguished, recapitulating the well-documented transcriptomic divergence between hindbrain and forebrain regions [65–67].

While end-repaired libraries generally aligned with their expected brain region, the displacement of A2AER, together with the concordant striatal placement of A2M2ER, signals degradation-driven bias rather than a regional identity mismatch. In contrast, because both C2 libraries were comparatively well-recovered (**Supplementary Table 8**), yet retained their cortical placement even under a marker-restricted projection, this is unlikely to reflect technical degradation, and thus it may instead indicate a cortical-derived transcriptomic signature in these libraries potentially arising from admixture of adjacent cortex tissue during dissection of the block prior to RNA extraction. In this way, our results show that decades-old FFPE brain RNA can retain region-discriminating protein-coding signatures, and that restricting to tissue-specific genes allows distinguishing degradation-driven misplacement from discordant transcriptional signatures possibly due to contamination of non-target nearby brain structures during sampling. Nevertheless, such clustering patterns might be affected by selection bias of marker genes included in our analyses, so further research is warranted to clarify brain regionality across FFPE samples.

Tissue specificity analysis using the tau index provided quantitative support for these observations. Of the 321 protein-coding genes and 14 microRNAs validated as tissue markers, many are canonical regional markers with well-characterized roles in cerebellar or striatal biology. Cerebellar vermis markers included the zinc finger transcription factors *ZIC1* and *ZIC4*, which define granule cell identity [68,69], the synaptic organizers *CBLN1* and *CBLN3*, essential for Purkinje cell synapse formation [70–72], the GABA-A receptor subunit *GABRA6*, exclusively expressed in cerebellar granule cells [73–75], and *PVALB*, a calcium-binding protein enriched in Purkinje cells [76]. In the corpus striatum, validated markers included the interneuron-associated neuropeptides *SST* and *NPY* [77,78], the neuron-specific kinase *CAMK2A* [79,80], and the calcium/calmodulin-dependent kinase *CAMKV* [81].

Besides, the functional enrichment of the two protein-coding marker sets was consistent with their regional origin. Cerebellar vermis markers were dominated by developmental and synapse-organization processes, matching the presence of developmental transcription factors and synaptic organizers among them (*ZIC1/4*, *CBLN1/3*, *RELN* or *PAX6*). In contrast, corpus striatum markers were instead dominated by active synaptic transmission and neurotransmitter release. Such divergence indicates that the region-specific expression retained in these FFPE brain samples carries coherent biological signal rather than unrelated isolated gene signatures. MicroRNA markers were similarly consistent with known regional biology. MiR-137, a robustly forebrain-enriched microRNA and a GWAS locus for schizophrenia [82,83], was among the strongest striatal markers, while the miR-124 family showed the expected cerebellar predominance [84–87]. The recovery of these well-established markers from tissue dating to the 1940s and 1950s demonstrates that regional molecular identity can survive the combined effects of post-mortem interval, formalin fixation, and prolonged storage.

Not all established markers met our validation criteria. Several canonical striatal genes, including *HTR2C* [88], *DRD2* [89], and *GNAL* [90], were detectable in FFPE corpus striatum samples but did not achieve sufficient transcriptional coverage for robust analysis. This highlights a key limitation of FFPE-based transcriptomics: while transcript presence and relative abundance can be captured, a comprehensive transcriptional reconstruction remains challenging for a subset of loci. Broader sampling across FFPE blocks and brain regions may improve transcript recovery, but further methodological advances will be needed to achieve more complete transcriptome profiling from highly degraded archival material.

### Implications for the Danish Brain Collection

Our results establish a practical framework for molecular profiling of the Danish Brain Collection and, by extension, similar historical FFPE archives. The combination of AllPrep extraction, T4 PNK end-repair, and single-stranded library preparation enabled recovery of tissue-specific expression profiles from samples up to 73 years old at the time of analysis. In parallel, a bioinformatic pipeline adapted from ancient and museum specimen studies [32,34] effectively accommodated the extensive fragmentation and chemical damage characteristic of these materials.

The Danish Brain Collection represents a unique resource for questions inaccessible to contemporary biobanks. These include the molecular correlates of historical diagnostic categories, the impact of changing environmental exposures and treatment regimes on brain molecular profiles, and retrospective investigation of infectious contributors to psychiatric disease in the pre-antibiotic and early antibiotic eras. The metagenomic and metatranscriptomic classification framework applied here, used primarily to assess contamination and library quality, provides a foundation for such investigations and could be extended to systematically screen for pathogen-derived nucleic acids across larger sample sets within the collection. The feasibility of recovering informative genomic and, critically, transcriptomic data from this pilot study opens the door to larger-scale investigations targeting additional defined diagnostic cohorts.

Nevertheless, several limitations warrant consideration: First, the sample size reported here is modest (four patients, 10 tissue blocks). Second, tissue representation is uneven across brain regions and individuals due to archival constraints, and the lack of matched fresh-frozen controls prevents direct quantification of information loss associated with FFPE processing. Third, DNA sequencing depth, while sufficient to assess mapping efficiency and damage patterns, would need to be increased beyond the target ∼10× coverage used here in order to support robust variant calling or comprehensive genomic analyses. This assessment, however, should be treated as a feasibility inference rather than a demonstrated result, as variant calling was not attempted in this pilot study. Regardless, an unknown number of FFPE blocks in the collection might not be suitable for genotyping irrespective of sequencing depth, as seen for patient D’s DNA data. Together, these observations indicate that DNA suitability for genotyping is likely to vary substantially across the collection depending on fixation chemistry, and should be evaluated on a per-sample basis rather than assumed uniformly across the whole archival material. In contrast, RNA from patient D showed mapping profiles broadly similar to the other donors despite the use of the acidic fixative, thus suggesting that RNA recovery may be less sensitive to fixation chemistry than DNA recovery. Such limitations can be addressed in future studies through expanded cohorts, deeper sequencing efforts, and the inclusion of matched contemporary controls where available.

## Conclusions

In summary, we demonstrate that decades-old FFPE brain tissue from the Danish Brain Collection is amenable to high-throughput DNA and RNA sequencing, retaining biologically meaningful, tissue-specific molecular signatures. These results establish the DBC as a viable resource for retrospective genomic and transcriptomic studies of neurological, neurodegenerative, and psychiatric disorders.

## Declarations

## Consent for publication

Not applicable

## Availability of data and materials

The sequence data generated in this study have been submitted to the NCBI BioProject database (https://www.ncbi.nlm.nih.gov/bioproject/) under accession numbers PRJNA1442830 and PRJNA1442847.

## Competing interests

The authors declare that they have no competing interests.

## Funding

EM-S acknowledges financial support from the Villum Fonden (Villum Experiment project no. 57875). NO is supported by the European Regional Development Fund project “Functional Omics Analysis of Metabolic Diseases to Advance Drug Discovery Research Excellence in LIOS - TARGETWISE” (no. 1.1.1.5/2/24/A/003). CK acknowledges financial support to this project through The Danish Association of Biomedical Laboratory Scientists.

## Authors’ contributions

EM-S and CK conceived this study and devised the experimental design. EM-S and CK secured funding. NO performed metagenomic and metatranscriptomic analyses. CRC and EHF performed laboratory work with support and supervision from SSTM and EM-S. CRC, EHF, EM-S, and NO analyzed the data, with input from CK and MTPG. CRC, EHF and EM-S visualized the data. MW facilitated ethics approval and sample acquisition. CRC, EHF, CK and EM-S wrote the manuscript with contributions and agreement from all authors.

## Acknowledgements

Not applicable

## Notes

### Competing Interest Statement

The authors have declared no competing interest.

### Summary of Updates

We have revised the manuscript according to Reviewer's requests.

