## Supplementary Figures for "Recovery of genomic and transcriptomic profiles from decades-old FFPE brain tissues"

^1^Department of Technology, Faculty of Health and Technology, University College Copenhagen, Copenhagen, Denmark. ^2^Center for Evolutionary Hologenomics, GLOBE Institute, University of Copenhagen. ^3^Department of Pathology, Rigshospitalet, The Capital Region of Denmark, Copenhagen, Denmark. ^4^Metabolic Research Group, Latvian Institute of Organic Synthesis, Riga, Latvia. ^5^University Museum, NTNU, Trondheim, Norway. ^6^Department of Pathology, Esbjerg Hospital, University Hospital of Southern Denmark, Esbjerg, Denmark. ^7^Department of Regional Health Research, University of Southern Denmark, Odense, Denmark. ^8^BRIDGE (Brain Research - Inter Disciplinary Guided Excellence), Department of Clinical Research, University of Southern Denmark, Odense, Denmark.

^*^Shared first authorship

^†^These authors contributed equally

***Corresponding authors:**

Emilio Mármol-Sánchez

Christina Kjær

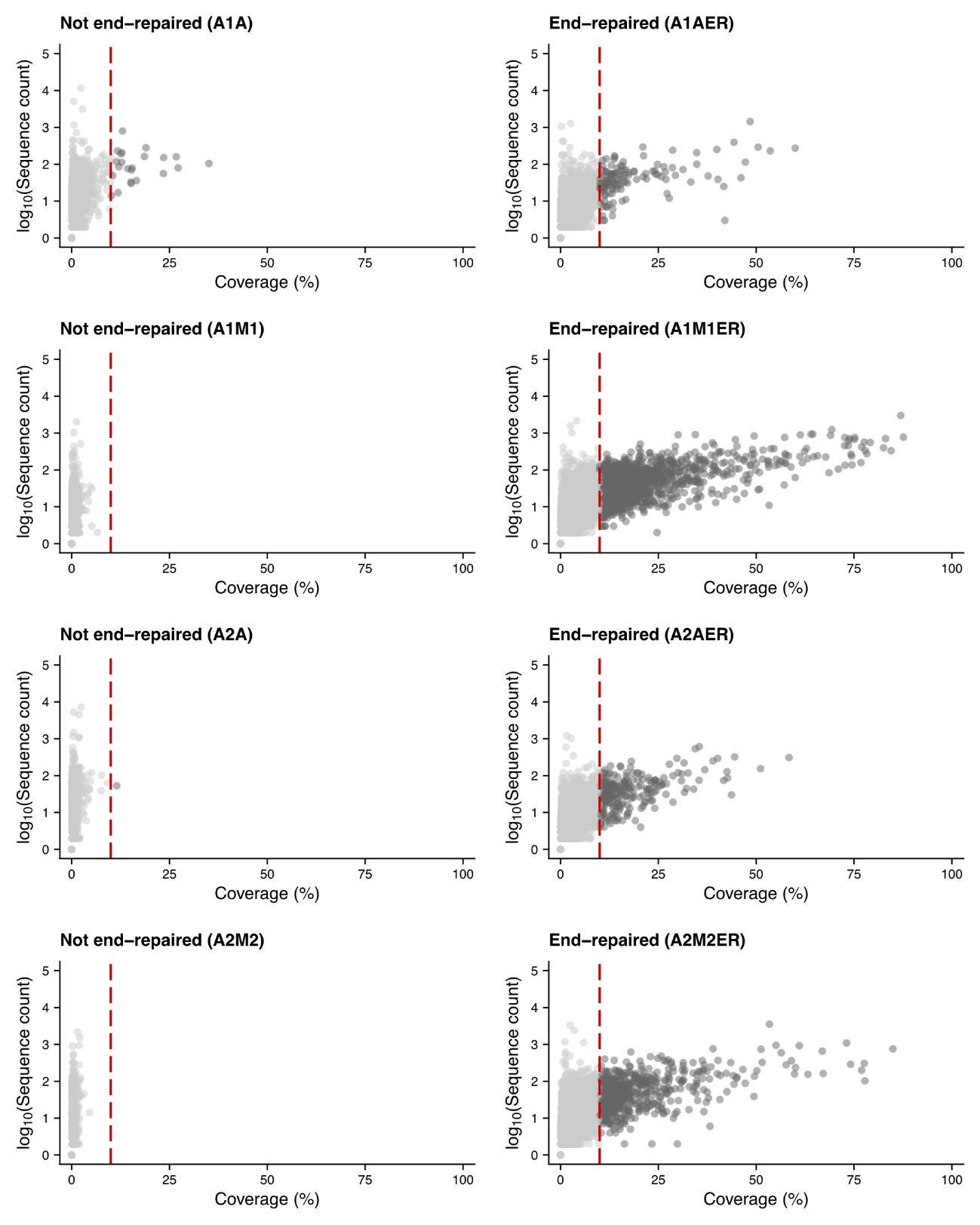

**Supplementary Figure 1: Per-sample sequence count vs. transcript coverage for protein-coding mRNA loci.** Comparison between libraries prepared from FFPE tissues with and without end-repair treatment in patient A. Each dot represents a protein-coding mRNA gene. The x-axis shows the percentage of the transcript length covered by mapped RNA fragments, while the y-axis shows the log_10_-transformed sequence count. The red dashed line marks a 10% coverage threshold, defined as the lower limit to consider any given protein-coding mRNA gene as successfully profiled.

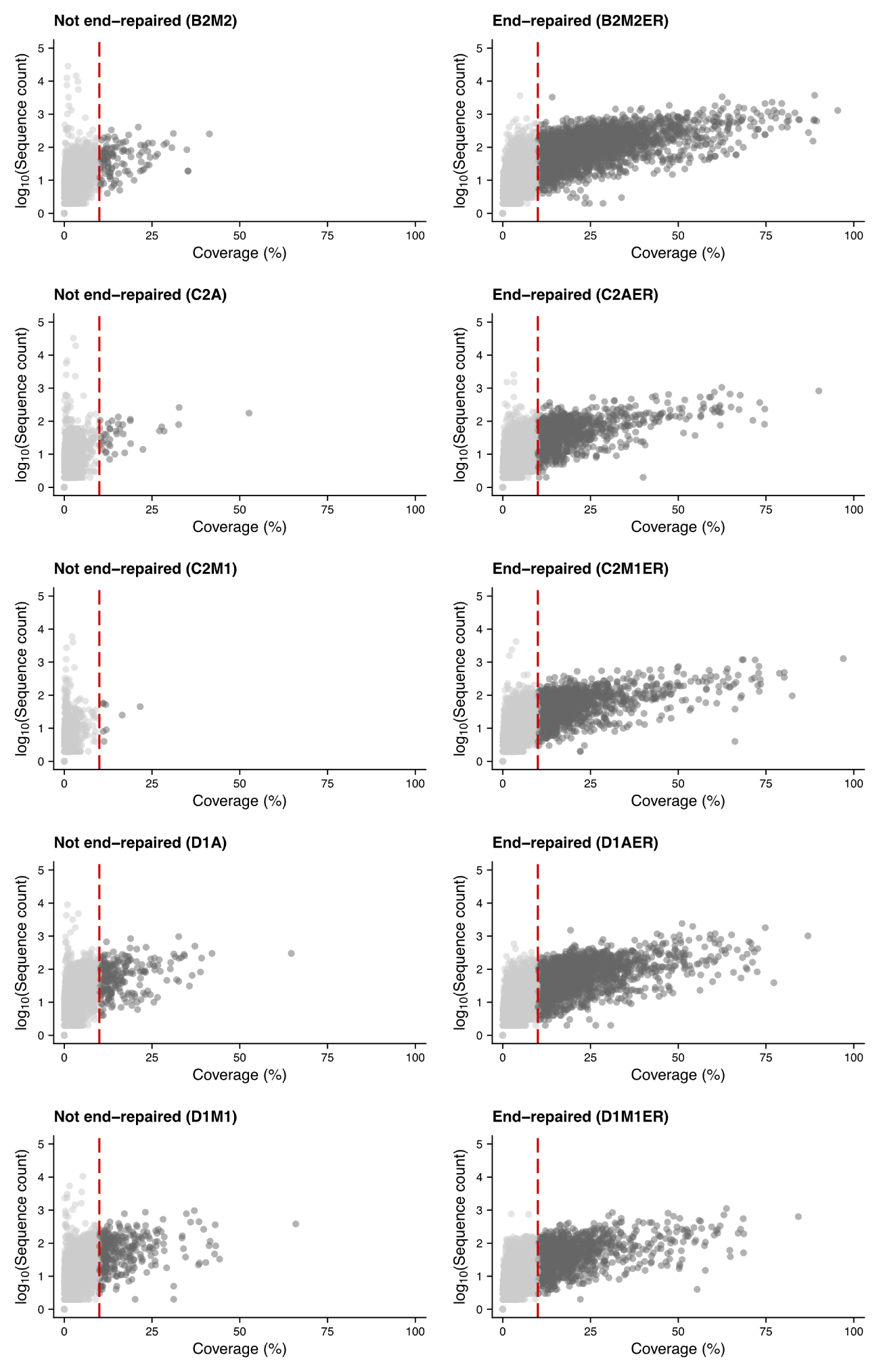

**Supplementary Figure 2: Per-sample sequence count vs. transcript coverage for protein-coding mRNA loci.** Comparison between libraries prepared from FFPE tissues with and without end-repair treatment in patients B, C & D. Each dot represents a protein-coding mRNA gene. The x-axis shows the percentage of the transcript length covered by mapped RNA fragments, while the y-axis shows the log_10_-transformed sequence count. The red dashed line marks a 10% coverage threshold, defined as the lower limit to consider any given protein-coding mRNA gene as successfully profiled.

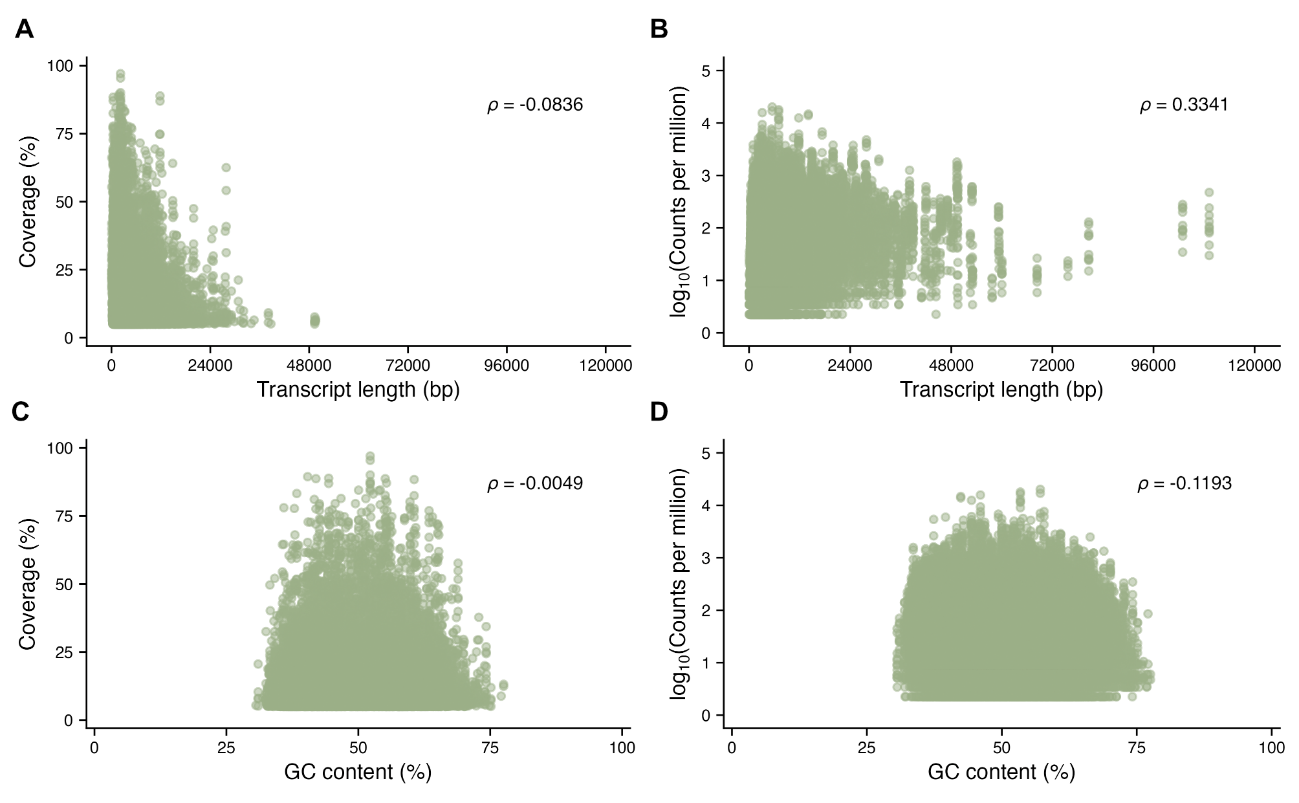

**Supplementary Figure 3: Association between gene features and FFPE mRNA recovery.** Per-gene transcript coverage (%) and abundance expresed in counts per million and log_10_-transformed plotted against transcript length (**A, B**) and transcript GC content (**C, D**), pooled across the ten end-repaired FFPE libraries. Only genes with transcript-wide coverage >10% and at least one log_10_ count per million are considered. Spearman’s rank correlation coefficient (*ρ*) between the two plotted variables is shown in each panel.

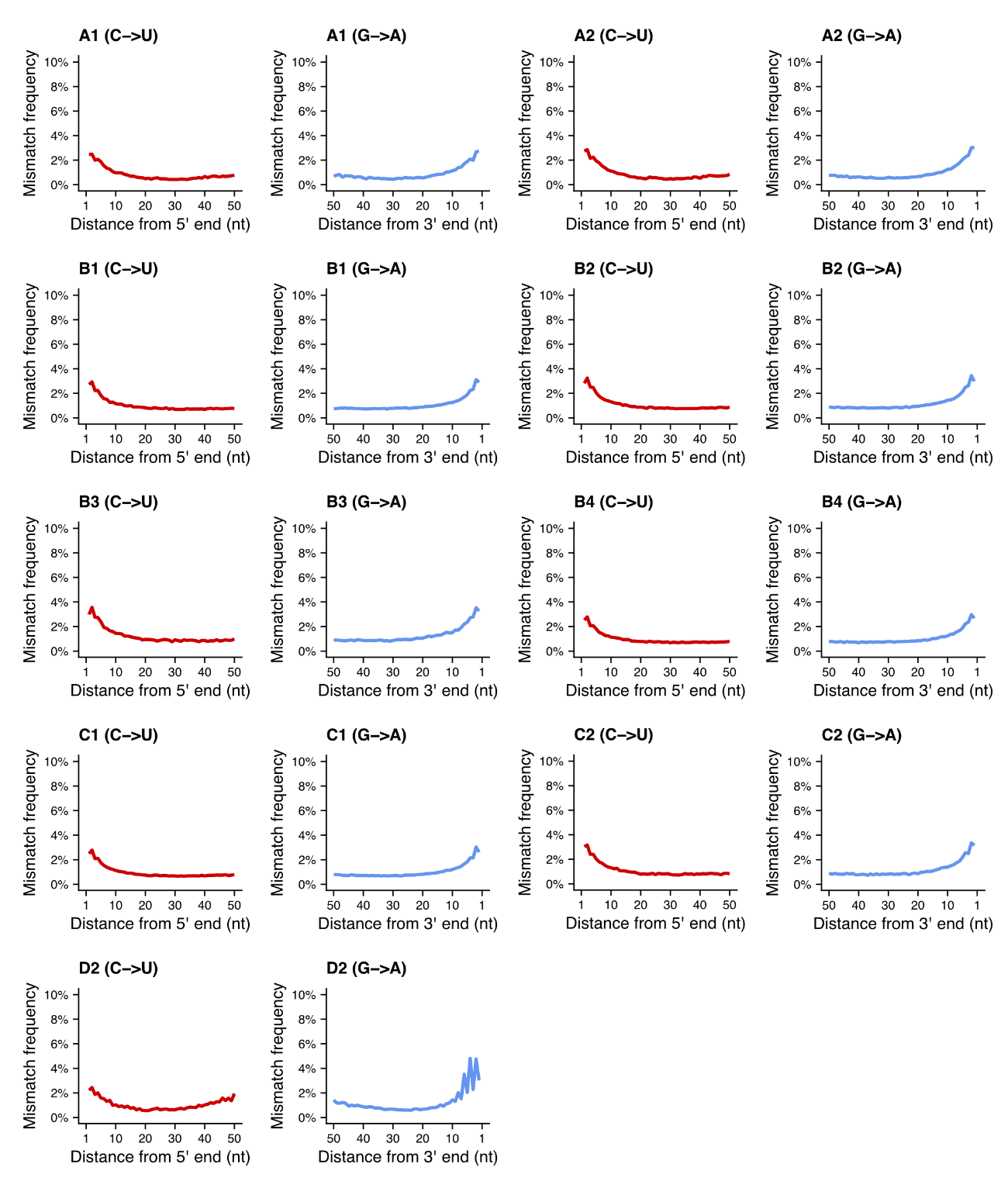

**Supplementary Figure 4: Cytosine deamination damage profiling of DNA sequencing data from FFPE brain samples.** G>A mismatches are considered as C>U deamination events for DNA sequences mapped to the negative strand. All 33 samples are included and damage profiles averaged according to individual and brain tissue (see **Table 1**).

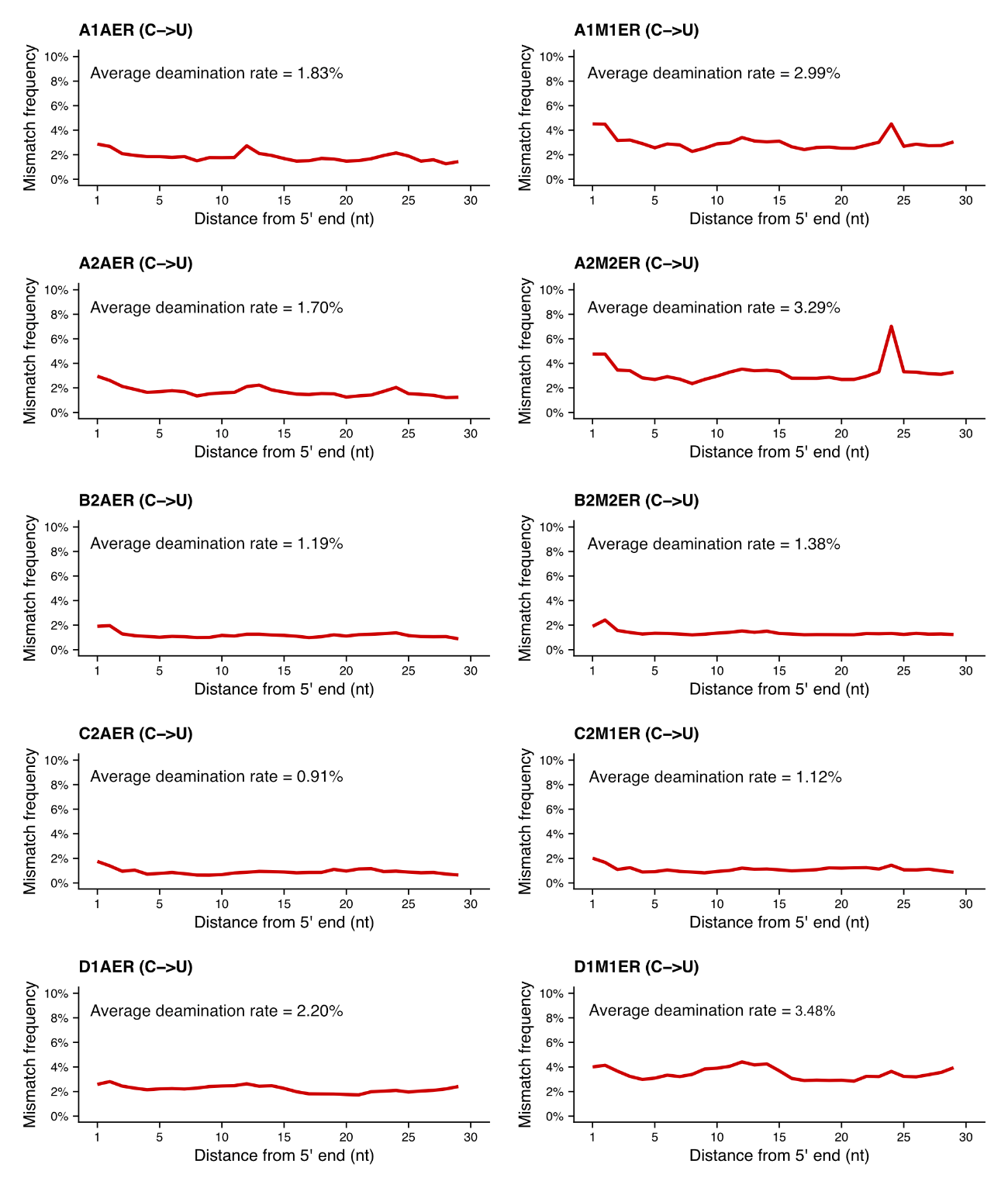

**Supplementary Figure 5:** **Cytosine deamination damage profiling of RNA sequencing data from FFPE brain samples after end-repair treatment.** The average C>U deamination rate is reported for each sample (n = 10).

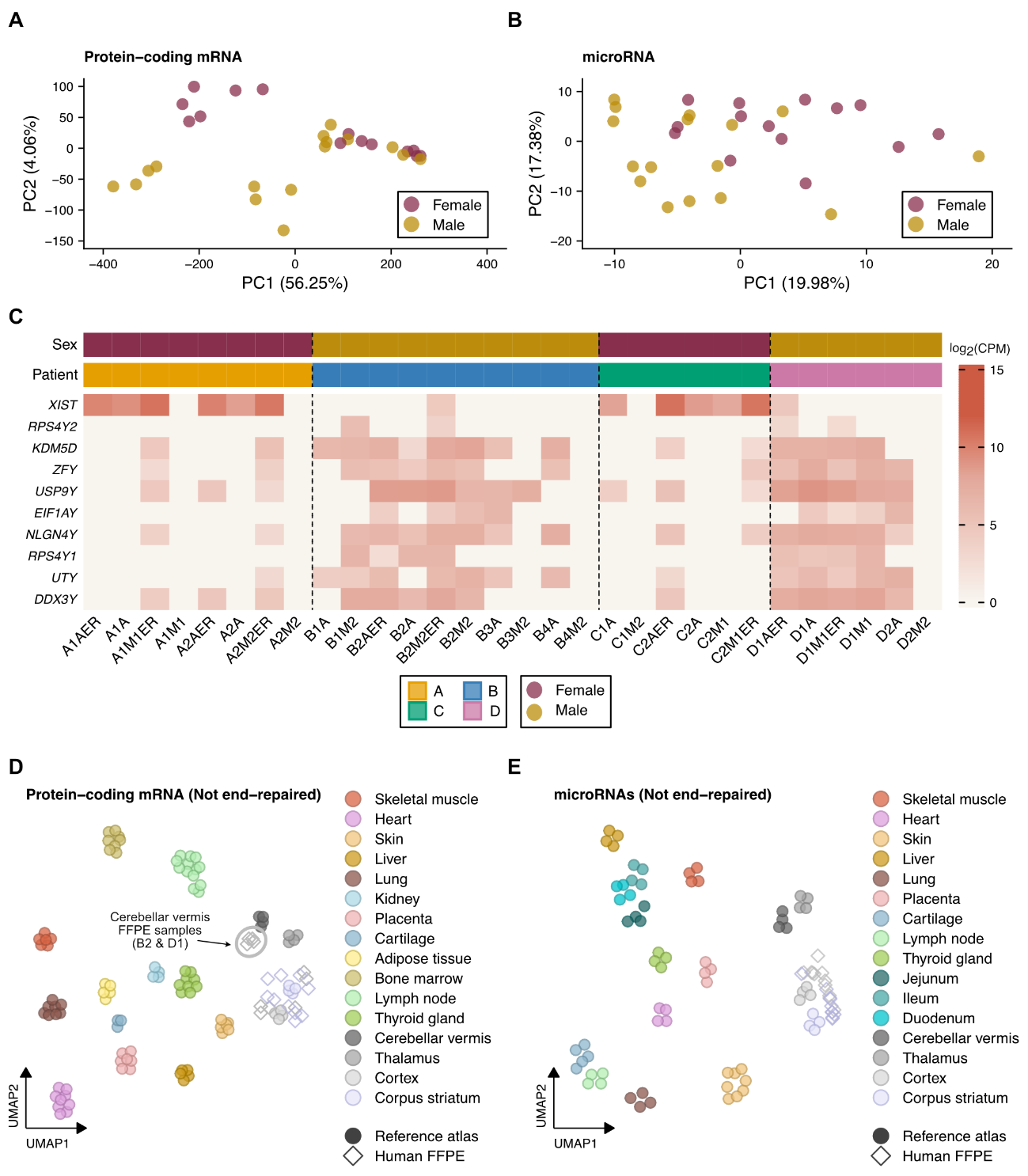

**Supplementary Figure 6: Sex-based PCA and heatmap, and multi-tissue UMAP projections for FFPE samples prepared without end-repair treatment.** **(A)** PCA of protein-coding mRNA expression across all 30 FFPE brain samples colored by sex. **(B)** PCA of microRNA expression across all 30 FFPE brain samples colored by sex. (**C**) Heatmap depicting sex-specific expression patterns of nine chromosome Y-linked marker genes and the lncRNA *XIST* across all RNA libraries from the four patients considered in this study. Intensity values for each gene are represented in log_2_-scaled counts per million (CPM) from normalized expression values using the trimmed mean of M-values (TMM) method. Each cell represents one individual RNA library. Dashed lines separate each of the four individual patients used in this pilot study. **(D)** UMAP embedding of protein-coding mRNA expression from FFPE brain samples without end-repair treatment projected onto a human multi-tissue RNA-seq reference atlas. **(E)** UMAP embedding of microRNA expression from FFPE brain samples without end-repair treatment projected onto a human multi-tissue small RNA-seq reference atlas.

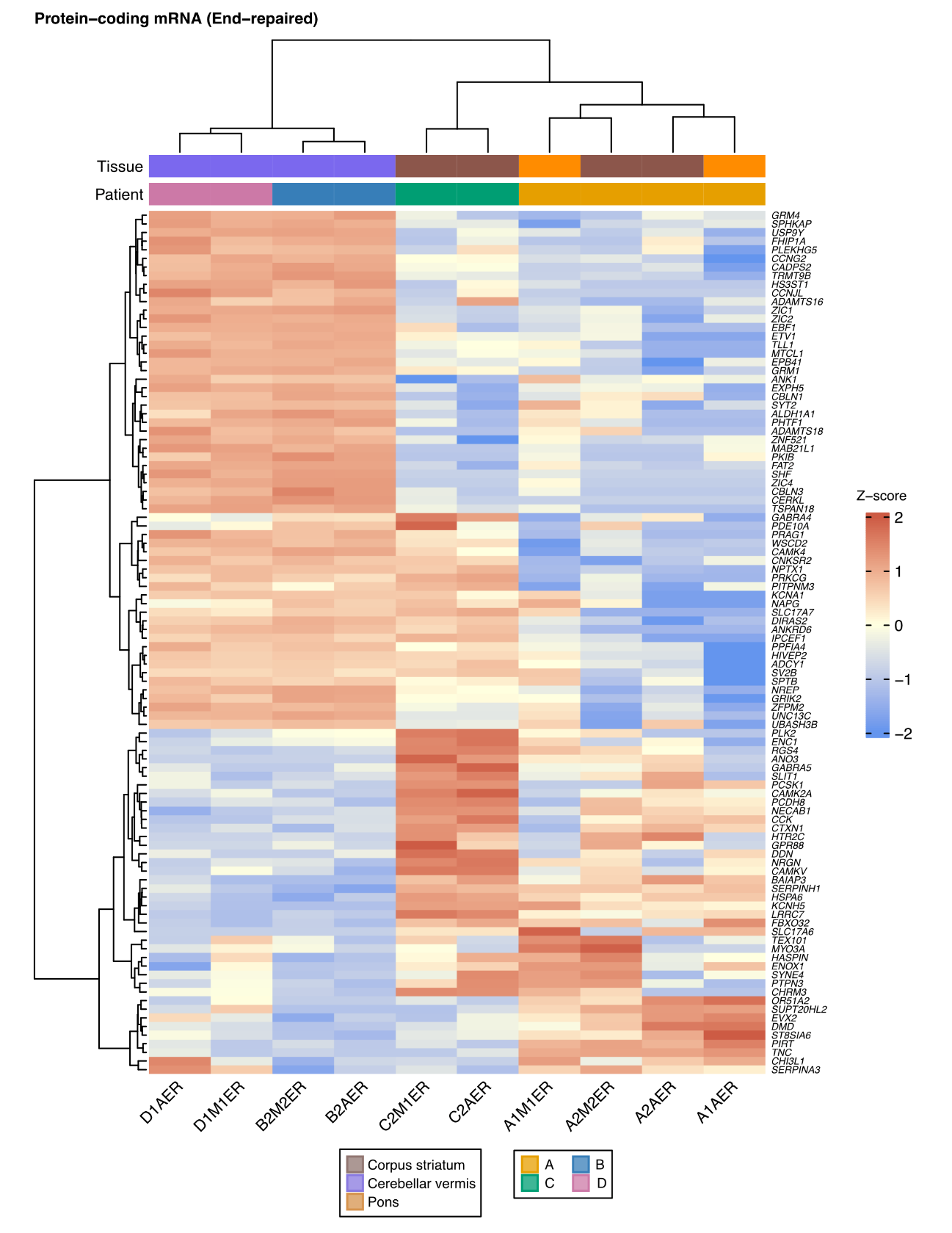

**Supplementary Figure 7: Hierarchical clustering of the top 100 most variable protein-coding mRNAs across end-repaired FFPE brain samples.** Expression values across samples (n = 10) are shown as row-wise z-scores computed on TMM-normalized log_2_ counts per million. Rows (genes) and columns (samples) are clustered using Ward's method with Euclidean distance.

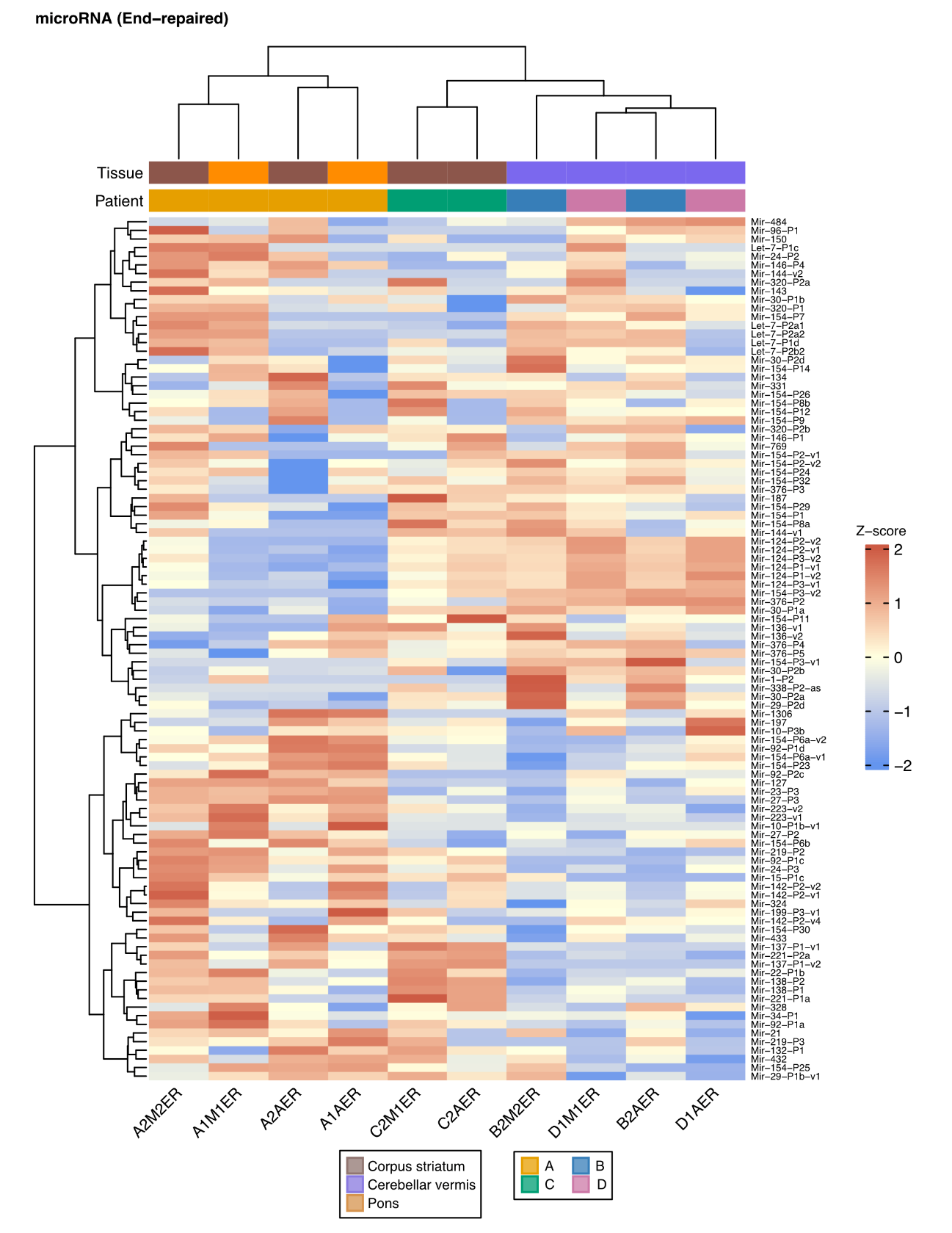

**Supplementary Figure 8: Hierarchical clustering of the top 100 most variable microRNAs across end-repaired FFPE samples.** Expression values across samples (n = 10) are shown as row-wise z-scores computed on TMM-normalized log_2_ counts per million. Rows (genes) and columns (samples) are clustered using Ward's method with Euclidean distance.
