## Supplementary Methods for "Recovery of genomic and transcriptomic profiles from decades-old FFPE brain tissues"

^1^Department of Technology, Faculty of Health and Technology, University College Copenhagen, Copenhagen, Denmark. ^2^Center for Evolutionary Hologenomics, GLOBE Institute, University of Copenhagen. ^3^Department of Pathology, Rigshospitalet, The Capital Region of Denmark, Copenhagen, Denmark. ^4^Metabolic Research Group, Latvian Institute of Organic Synthesis, Riga, Latvia. ^5^University Museum, NTNU, Trondheim, Norway. ^6^Department of Pathology, Esbjerg Hospital, University Hospital of Southern Denmark, Esbjerg, Denmark. ^7^Department of Regional Health Research, University of Southern Denmark, Odense, Denmark. ^8^BRIDGE (Brain Research - Inter Disciplinary Guided Excellence), Department of Clinical Research, University of Southern Denmark, Odense, Denmark.

^*^Shared first authorship

^†^These authors contributed equally

***Corresponding authors:**

Emilio Mármol-Sánchez

Christina Kjær

**Supplementary Methods**

**DNA-only extraction**

DNA was extracted in a first batch from ten independent FFPE blocks (see **Table 1** and **Supplementary Table 1**) using the DNeasy Blood & Tissue Kit (Qiagen, Germany), and following the manufacturer’s instructions with the following modifications: Approximately 25 mg of tissue was scraped from each FFPE block using single-use sterile scalpels and tweezers, and then rinsed twice for 1 min. each with 1× PBS by repeated vortexing and gentle shaking of the tubes. The PBS solution was then removed and the tissue piece was chopped into smaller pieces prior to the addition of 180 µL of ATL buffer. The mixture was then heated to 90°C for 10 min. to allow decrosslinking [1], and then left to cool down at room temperature for 5 min. A total of 40 µL of Proteinase K solution from the DNeasy Blood & Tissue Kit (Qiagen) were next added and incubated at 56°C overnight for digestion. Following steps were according to the manufacturer’s protocol. Final elution was done in 50 µL AE buffer and stored at -20°C until further use. A second batch of DNA extraction was performed from five of the initial FFPE blocks (B1, B2, B4, C1 and D2, see **Supplementary Table 1**) totaling 10 samples (n = 5 Melted, and n = 5 original FFPE) using the DNeasy Blood & Tissue Kit (Qiagen, Germany) for a total of 10 extracts to compare the effect of the paraffin melting treatment in DNA yield and sequence fragment length.

Six additional independent extractions were carried out in a third batch from samples B1, B2 and B4 on the original FFPE (n = 3) and the melted (n = 3) block fragments (**Supplementary Table 1**), using the DNA extraction DNeasy Blood & Tissue Kit as before, but with the prior addition of a deparaffinization step as follows: after the tissue had initially been cut into smaller pieces, 320 µL of Deparaffinization Solution (Qiagen, Germany) was added, and samples were incubated at 56°C for 3 min., prior to the addition of 180 µL of ATL buffer. Following steps for overnight digestion were as described above, after which the lower phase was carefully transferred to clean Eppendorf tubes and processed according to the remainder of the manufacturer’s protocol for DNA extraction.

**RNA-only extraction**

RNA was extracted from the ten original FFPE brain tissue samples (A1, A2, B1, B2, B3, B4, C1, C2, D1 and D2, see **Table 1** and **Supplementary Table 1**) using the mirVana^TM^ miRNA Isolation Kit (Thermo Fisher Scientific). Additional pretreatment steps to optimize recovery from FFPE material were implemented as follows: Approximately 30 mg of FFPE tissue was excised from each block with single-use sterile scalpels and tweezers and the excess paraffin was manually trimmed. Samples were then incubated at 58°C for 10 min. to melt the excess paraffin, washed thrice with 500 µL of 1× PBS and heat-treated at 75°C for 15 min. to facilitate decrosslinking before RNA isolation. A total of 1 mL Lysis/Binding buffer was then added, and tissue homogenization was performed using Lysis Matrix M 2 mL tubes (MP Biomedicals) and a FastPrep-24 5G bead-beating grinder and lysis system using the preprogrammed Mouse Brain setting (6 m/s for 40 s.), repeated thrice with tube cooling in cold racks between each cycle to prevent overheating. Subsequent steps were performed according to the manufacturer’s protocol for RNA isolation with acidic phenol:chloroform solution (Thermo Fisher Scientific). The final elution of the RNA extract in 23 µL of nuclease-free water was repeated twice, and finally stored at -70°C until further use.

**Simultaneous DNA and RNA extraction**

A fourth batch of dual DNA & RNA extraction from the ten original FFPE samples was performed using the AllPrep DNA/RNA FFPE Kit (Qiagen) according to the manufacturer’s instructions (**Table 1** and **Supplementary Tables 1-2**), with minor modifications as follows: For RNA purification, total RNA including the small RNA fractions was carried out by first shaving thin layers of tissue (~10 mg) cut directly from the original FFPE blocks with single-use sterile scalpels and tweezers, and excess solid paraffin was trimmed. The shavings were then treated with 320 µL Deparaffinization Solution (Qiagen, Germany). Following this, the rest of the extraction was carried out according to the manufacturer’s protocol. Following silica column-based nucleic acid purification, DNA molecules were eluted in 50 µL ATE buffer. Finally, RNA molecules were eluted in 23 µL nuclease-free water and stored at -70°C until further use.

**DNA library preparation**

Prior to library preparation, each DNA extract was diluted to equimolar concentration in a volume of 32 µL using EB buffer (see **Supplementary Table 1** for further details). Library preparation was then performed in three sequential steps comprising end-repair, adapter ligation and fill-in reaction. End-repair was first performed by mixing 23 µL of each DNA dilution and 8 µL mastermix consisting of the following components: 4.0 µL 10× T4 DNA ligase buffer (NEB), 2.2 µL Reaction booster enhancer, 0.4 µL T4 DNA polymerase (NEB), 1 µL T4 Polynucleotide Kinase (NEB), and 0.4 µL of 25 mM dNTP (GeneON) per sample. The end-repair reaction was then incubated for 30 min. at 20°C followed by 30 min. at 65°C in a prewarmed thermocycler. For adapter ligation, 2 µL of P5+P7 adapter solution (10 µM or 20 µM, 10-50:1 adapter:insert ratio, final concentration >0.2 µM) was added to the reaction tubes and mixed by pipetting, followed by the addition of 8 µL mastermix consisting of 6 µL 50% PEG-4000 (Sigma-Aldrich), 1 µL 10× T4 DNA ligase buffer (NEB) and 1 µL T4 DNA ligase (NEB). The ligation reaction was then incubated for 30 min. at 20°C followed by 10 min. at 65°C, and cooled to 4°C in a prewarmed thermocycler. The fill-in reaction was performed by adding 10 µL of mastermix to each sample consisting of 5.6 µL nuclease-free water, 2.0 µL 10× Isothermal amplification buffer (NEB), 1.6 µL Bst 2.0 Warmstart Polymerase (NEB, 8 U/µL) and 0.8 µL dNTP (25mM), resulting in a final reaction volume of 60 µL for each sample. The reaction was incubated for 15 min. at 65°C followed by 15 min. at 80°C, and finally cooled to 4°C in a prewarmed thermocycler. Five negative controls were included: three with EB buffer instead of DNA template and one of each with AE buffer and ATE buffer as input.

Library clean-up was then performed using 100 µL HighPrep PCR beads (MagBio Genomics). The beads were first mixed with the DNA library and incubated for 5 min. at room temperature, before being placed on a magnetic rack for 3 min. The supernatant was discarded, and the beads were then washed twice with 200 µL freshly prepared 80% ethanol. After this, the beads were left to air-dry for 3 min. The cleaned DNA library was finally eluted in 30 µL EBT buffer and incubated at 37°C for 10 min., then placed on the magnetic rack, and the DNA library eluate transferred to a new tube and stored at -20°C until further use.

Quantitative polymerase chain reaction (qPCR) was performed using 1 µL of 1:10 dilutions of the DNA libraries and 19 µL of mastermix consisting of 12.88 µL nuclease-free water, 2 µL 10× PCR Gold Buffer (Thermo Fisher Scientific), 2 µL MgCl₂ Solution 25 mM (Thermo Fisher Scientific), 0.8 µL SYBR Green, 0.2 µL Recombinant albumin 20mg/mL (NEB), 0.16 µL 25 mM dNTP (GeneON), 0.16 µL AmpliTaq Gold DNA Polymerase 5 U/µL (Thermo Fisher Scientific), 0.4 µL 10 µM P5 qPCR primer and 0.4 µL 10 µM P7 qPCR primer per reaction. A positive control and corresponding negative controls consisting of nuclease-free water as input instead of DNA library were included to verify assay performance and check for contamination. The qPCR was performed on a Mx3005P qPCR system (Agilent Technologies) with the following program: 95°C for 10 min., followed by 40 cycles of 95°C for 30 s., 60°C for 1 min. and 72°C for 1 min. A dissociation curve was generated by applying the following program: 95°C for 1 min., 55°C for 30 s., 95°C for 30 s.

The PCR indexing was performed by adding 38 µL mastermix to 10 µL of DNA library for each sample. The mastermix consisted of 25.5 µL nuclease-free water, 5 µL 10× PCR Gold Buffer (Thermo Fisher Scientific), 5 µL MgCl₂ Solution 25 mM (Thermo Fisher Scientific), 0.5 µL 25 mM dNTP (GeneON), 1 µL Recombinant albumin 20mg/mL (NEB) and 1 µL AmpliTaq Gold DNA Polymerase 5 U/µL (Thermo Fisher Scientific) per reaction. Libraries were then indexed using 1 µL 10 µM P5 and P7 8-bp dual index primers (unique for each library). A negative control containing nuclease-free water instead of DNA was included. Libraries were then amplified for sequencing in an Applied Biosystems 2720 Thermal Cycler (Thermo Fisher Scientific) using the following program: 95°C for 10 min., followed by a number of cycles of 95°C for 30 s., 60°C for 1 min. and 72°C for 1 min., followed by 5 min. at 72°C. Each library was assigned to a specific number of PCR cycles based on the qPCR, where the number of cycles were calculated based on the middle of the amplification curve (exponential fraction) for each sample. Each indexed library was finally purified with 60 µL HighPrep PCR beads (MagBio Genomics) as described above and eluted in 30 µL EBT. The final DNA libraries were then stored at -20°C until further use.

**End-repair**

End-repair was carried out on 10 RNA extracts (RNA extracted with either the mirVana or the AllPrep kits from samples A1, A2, B2, C2, D1, see **Supplementary Table 2**) before the start of the protocol for the RNA library preparation using T4 Polynucleotide Kinase (NEB), T4 Polynucleotide Kinase Reaction Buffer 10× (NEB) and Adenosine 5'-triphosphate (NEB). Briefly, a total of 5 µL of RNA extract was mixed with 2 µL ATP, 1 µL T4 PNK, and 2 µL T4 PNK Reaction Buffer. The reaction was incubated at 37°C for 30 min., followed by 65°C for 20 min., then held at 4°C until further processing. Finally, 5 µL of the incubation product for each sample was used as input for RNA library preparation. The miRNA Control included in the NEXTFLEX Small RNA-Seq Kit was used as a positive control sample. Two negative controls were included: one where nuclease-free water (NFW) was used for the end-repair step instead of RNA and treated like the other samples (Neg NFW ER) and one where NFW was used as the input instead of end-repaired RNA (Neg NFW).

**RNA library preparation**

cDNA libraries from RNA extracts were prepared using a total of 20 samples (RNA extracted with the mirVana or the AllPrep kits from the 10 original FFPE samples) with the NEXTFLEX Small RNA-Seq Kit v4 including Unique Dual Indexes (UDIs) (Revvity), without size selection, and according to the manufacturer’s instructions (**Supplementary Table 2**).

The NEXTFLEX 3' Adenylated adapter and NEXTFLEX 5' Adapter were first diluted 1:4 with nuclease-free water, according to the manufacturer’s recommendations for poor-quality samples. The miRNA Control included in the NEXTFLEX Small RNA-Seq Kit was used as a positive control sample. A negative control with nuclease-free water as input instead of RNA extract was included. A negative PCR control was also included, where nuclease-free water was used instead of PCR mastermix.

**Metagenomics & metatranscriptomics**

Taxonomic classification of DNA and RNA sequences was performed to characterize the species composition of each library, including endogenous human content and any exogenous microbial contamination. The analysis was carried out on adapter-trimmed sequences without PCR deduplication, retaining only sequences of at least 30 nucleotides. DNA and RNA fragments were classified using KrakenUniq v0.7.3 [2] with a custom database [3] built from the microbial part of the NCBI non-redundant nucleotide (NT) database (<https://doi.org/10.17044/scilifelab.20518251>). Species were reported as detected (**Supplementary Tables 9-10**) if they met minimum thresholds of ≥500 unique k-mers and ≥50 assigned sequences.

**Clustering parameters**

UMAP algorithm initialization and embedding were computed for both protein-coding and miRNA quantification data independently, and defined with the following parameters: n_neighbors = 5, metric = "pearson", spread = 1, random_state = 40. Normalized FFPE RNA data from samples with and without end-repair treatment were then projected onto the learned UMAP embedding using the predict function in R v4.4.2. For brain-specific clustering, two UMAP embeddings were generated: one using all annotated protein-coding genes (**Fig. 4A**), and one restricted to tissue-specific genes, defined as genes with τ ≥ 0.5 and peak expression in either cortex, cerebellar vermis, or corpus striatum (**Supplementary Table 19**; **Fig. 4B**). Because these protein-coding genes were selected based on the previous gene expression atlas (E-MTAB-2836 plus additional manually inserted samples, **Supplementary Table 3**), and independently of the GTEx reference (**Supplementary Table 4**) used for the embedding, this projection is not biased by the composition of the reference being tested.

**Gene feature association analyses**

For each protein-coding gene, transcript length and GC content (%) were obtained from the reference transcriptome annotation considering the longest transcript per gene. Transcript coverage (percentage of transcript body covered by mapped sequences) and abundance expressed in counts per million (CPM) and log_10_-transformed (adding a pseudo-count of 1) were computed and pooled across the ten end-repaired FFPE libraries. Genes were considered only if they showed reliably detectable recovery (coverage >10% and at least 1 mapped sequence expressed in CPM and log_10_-transformed). Associations between gene features (transcript length, and transcript GC content) and recovery metrics (coverage, abundance) were quantified using Spearman’s rank correlation coefficient (*ρ*).
